# Puffaligner: An Efficient and Accurate Aligner Based on the Pufferfish Index

**DOI:** 10.1101/2020.08.11.246892

**Authors:** Fatemeh Almodaresi, Mohsen Zakeri, Rob Patro

## Abstract

**Motivation:** Sequence alignment is one of the first steps in many modern genomic analyses, such as variant detection, transcript abundance estimation and metagenomic profiling. Unfortunately, it is often a computationally expensive procedure. As the quantity of data and wealth of different assays and applications continue to grow, the need for accurate and fast alignment tools persists.

**Results:** In this paper, we introduce PuffAligner, a fast, accurate and versatile aligner built on top of the Pufferfish index. PuffAligner is able to produce highly-sensitive alignments, similar to those of Bowtie2, but much more quickly. While exhibiting similar speed to the ultrafast STAR aligner, PuffAligner requires considerably less memory to construct its index and align reads. PuffAligner strikes a desirable balance with respect to the time, space, and accuracy tradeoffs made by different alignment tools, and provides a promising foundation on which to test new alignment ideas over large collections of sequences.

**Availability:** PuffAligner is a free and open-source software. It is implemented in C++14 and can be obtained from https://github.com/COMBINE-lab/pufferfish/tree/cigar-strings

## 1 Introduction

Since its introduction, next generation sequencing (NGS) has been widely used as a low-cost and accessible technology to produce high-throughput sequencing reads for many important biological assays. The sequencing data that is generated in the form of short reads, drawn from longer molecular fragments, and finding the optimal alignments of these short reads to some reference is a necessary first step for many downstream biological analyses. The process of finding the segment on the reference that is most similar to the query read, and therefore most likely to be the source of the fragment from which the read was drawn, is known as read mapping or read alignment.

The main goal in read alignment is to find alignments of contiguous sub-string of the underlying reference that yields a minimum edit distance (or maximum alignment score) between the read and the reference sequence at the alignment position. If the reads are paired-end, characteristics other than the alignment score can be used to filter spurious alignment locations, such as orientation of each end of the alignment pair (forward or reverse) or distance between the alignments corresponding to reads that are ends of the same fragment.

Short-read aligners are a major workhorse of modern genomics. Given the importance of the alignment problem, a tremendous number of different tools have been developed to tackle this problem. Some widely used examples are BWA^1^, Bowtie2^2^, Hisat2^3,4^ and STAR^5^. Existing alignment tools use a variety of indexing methods. Some tools, such as BWA, Bowtie2, and STAR use a full-text index over the reference sequences; BWA and Bowtie2 use variants of the FM-index, while STAR uses a suffix array.

A popular alternative approach to full-text indices is to instead, index sub-strings of length *k* (*k*-mers) from the reference sequence. Trading off index size for potential sensitivity, such indices can either index all of the *k*-mers present in the underlying reference, or some uniform or intelligently-chosen sampling of *k*-mers. There are a large variety of *k*-mer -based aligners, including tools like the Subread aligner^6^, SHRiMP2^7^, mrfast^8^, and mrsfast^9^. To reduce the index size, one can choose to select specific *k*-mers based on a winnowing (or minimizer) scheme. This approach has been particularly common in tools designed for long-read sequence alignment like mashmap^10^ and minimap2^11^.

Recently, a set of new indices for storing *k*-mers have been proposed based on graphs, specifically de Bruijn graphs (dBg). A de Bruijn graph is a graph over a set of distinct *k*-mers where each edge connects two neighboring *k*-mers that appear consequently in a reference sequence and therefore, overlap on “*k* − 1” bases. Kallisto^12^, deBGA^13^, BGreat^14^, BrownieAligner^15^, and Pufferfish^16^ are some tools which use an index constructed over the de Bruijn graph built from the reference sequences. Cortex^17^, Vari^18^, rainbowfish^19^, and mantis^20^ are also tools that use a colored compacted de Bruijn graph for building their index over a set of raw experiments. All these approaches cover a wide range of the possible design space, and different design decisions yield different performance tradeoffs.

Generally, the fastest aligners (like STAR) have very large memory requirements for indexing, and make some sacrifices in sensitivity to obtain their speed. On the other hand, the most sensitive aligners (like Bowtie2) have very moderate memory requirements, but obtain their sensitivity at the cost of a higher runtime. Maintaining the balance between time and memory is especially more critical while aligning to a large set of references, like a large collection of microbial and viral genomes which may be used as an index in microbiome or metagenomic studies. As both the collection of reference genomes and the amount of sequencing data grows quickly, it is import for alignment tools to achieve a time-space balance without loosing sensitivity.

Based on the compact Pufferfish^16^ index, we introduce a new aligner called PuffAligner, that we believe strikes an interesting and useful balance in this design space. PuffAligner is designed to be a highly-sensitive alignment tool while, simultaneously, placing a premium on computational overhead. By using the colored compacted de Bruijn graph to factor out repeated sub-sequences in the reference, it is able to leverage the speed and cache friendliness of hash-table based aligners while still controlling the growth in the size of the index; especially in the context of redundant reference sequences. By carefully exploring the alignment challenges that arise in different assays, including single-organism DNA-seq, RNA-seq alignment to the transcriptome, and metagenomic sequencing, we have engineered a versatile tool that strikes desirable balance between accuracy, memory requirements and speed. We compare PuffAligner to some other popular aligners and show how it navigates these different tradeoffs.

## 2 Results

For measuring the performance of PuffAligner and comparing it to other aligners, we have designed a series of experiments using both simulated and experimental data from different sequencing assays. We compare PuffAligner with Bowtie2^2^, STAR^5^ and deBGA^13^. Bowtie2 is a popular, sensitive and accurate aligner with the benefit of having very modest memory requirements. STAR requires a much larger amount of memory, but is much faster than Bowtie2 and can also perform “spliced alignment” against a reference (which PuffAligner, Bowtie2, and deBGA currently do not allow). deBGA, is most-related tool to PuffAligner conceptually, as it is an aligner with a colored compacted de Bruijn graph-based index that is focused on exploiting redundancy in the reference sequence.

We use different metrics to assess both the performance and accuracy of each method on a variety of types of sequencing samples. These experiments are designed to cover a variety of different use-cases for an aligner, spanning the gamut from situations where most alignments are expected to be unique (DNA-seq), to situations where each fragment is expected to align to many loci with similar quality (RNA-seq and metagenomic sequencing), and spanning the range of index sizes from small transcriptomes to large collections of genomes.

First, we show PuffAligner exhibits similar accuracy for aligning DNA-seq reads to Bowtie2, but it is considerably faster. In the case of experimental reads, since the true origin of the read is unknown, we use measures such as mapping rate and concordance of alignments to compare the methods. Furthermore, we evaluate the accuracy of aligners by aligning simulated DNA-seq reads that include variation (single-nucleotide variants and small indels with respect to the reference). For aligning RNA-seq reads, we compare the impact of alignments produced by each aligner on downstream analysis such as abundance estimatation. Finally, we show PuffAligner is very efficient for aligning metagenomic samples where there is a high degree of shared sequence among the reference genomes being indexed. We also illustrate that using alignments produced by PuffAligner yields the highest accuracy for abundance estimation of metagenomic samples.

### 2.1 Configurations of aligners in the experiments

The performance of each tool is impacted by the different alignment scoring schemes they use, e.g. different penalties for mismatches, and indels. To enable a fair comparison, we attempted to configure the tools so as to minimize divergences that simply result from differences in the scoring schemes. For the experiments in this paper, we use Bowtie2 in a near-default configuration (though ignoring quality values), and attempt to configure the other tools, as best as possible, to operate in a similar manner.

The deBGA scoring scheme is not configurable, so we use this aligner in the default mode (unfortunately, the inability to disable local alignment and forcing just computation of end-to-end alignments in deBGA makes certain comparisons particularly difficult). For PuffAligner we use a scheme as close to Bowtie2 as possible. The maximum possible score for a valid alignment in Bowtie2 is 0 (in end-to-end mode) and each mismatch or gap subtracts from this score. Bowtie2 uses an affine gap penalty scoring scheme, where opening and extending a gap (insertion or deletion) have a cost of 5 and 3 respectively. For DNA-seq reads, we configure STAR to allow as many mismatches as Bowtie2 and PuffAligner by setting the options “`--outFilterMismatchNoverReadLmax 0.12`” and “`--outFilterMismatchNmax 1000`”. Also, we use the option “`--alignIntronMax 1`” in STAR to perform non-spliced alignments while aligning genomic reads. For RNA-seq reads, STAR has a set of parameters which we change in our result evaluations, and which are detailed below in the relevant sections.

In Bowtie2 we also use the option `--gbar 1` to allow gaps anywhere on the read except within the first nucleotide (as the other tools have no constraints on where indels may occur). Furthermore, for consistency, we also run Bowtie2 with the option “`--ignore-quals`”, since the other tools do not utilize base qualities when computing alignment scores.

As explained in Section 4.1, for the sake of performance, highly repeated anchors (more than a user-defined limit) will be discarded before the alignment phase. This threshold is by default equal to 1000 in PuffAligner. We set the threshold to the same value for STAR and deBGA using options `--outFilterMultimapNmax 1000` and `-n 1000` respectively. There is no such option exposed directly in Bowtie2.

Since PuffAligner finds end-to-end alignments for the reads, we are also running other tools in end-to-end mode, which is the default alignment mode in Bowtie2 as well. In STAR we enable this mode using the option `--alignEndsType EndToEnd`. In the case of deBGA, although the documentation suggests it is *not* supposed to find local alignments by default, the output SAM file contains many reads with relatively long soft clipped ends, so if a read is not aligned end-to-end, deBGA reports the local alignment for that. We were not able to find any option to force deBGA to perform end-to-end alignments for all reads, and so we have compared it in the configuration in which we were able to run it.

For aligning DNA-seq samples, each aligner is configured to report a single alignment, which is the primary alignment, for each read. Bowtie2 outputs one alignment per read by default. To replicate this in the other tools, we use the option `--outSAMmultNmax 1` in STAR, `-o 1 -x 1` in deBGA, and `--primaryAlignment` in PuffAligner.

### 2.2 Alignment of whole genome sequencing reads

First, we evaluate the performance of PuffAligner with a whole genome sequencing (WGS) sample from the 1000 Genomes project^21^.We downloaded the `ERR013103` reads from sample `HG00190`, which is a low-coverage sample from a Finnish male, sequenced in Finland.^*^. There are 18,297,585 paired-end reads, each of length 108 nucleotides in this sample. Using fastp^22^, we remove low quality ends and adapter sequences from these reads. After trimming, there are 15,404,412 reads remaining in the sample. Indices for each of the tools are built over all DNA chromosomes of the latest release of the human genome (v33) by gencode^†23^.

In this experiment, all aligners are configured report only concordant alignments, i.e., only pairs of alignments that are cocordant and within the “maximum fragment length” shall be reported. The maximum fragment length in all aligners is set to 1000, using the option `--alignMatesGapMax 1000` in STAR, `--maxins 1000` in Bowtie2 and `-u 1000 -f 0` in deBGA. The default value for the maximum fragment length in PuffAligner is set to 1000, the user can cofigure this value by using the flag `--maxFragmentLength`. This concordance requirements also prevents Bowtie2, PuffAligner, and STAR from aligning both ends of a paired end read to the same strand.

The alignment rate, run-time memory usage and running time for all the aligners are presented in 1. The reason that deBGA has the highest mapping rate in 1 compared to other tools is that it is local alignments for the reads that are not alignable end-to-end under the scoring parameters for the other tools. Bowtie2 and PuffAligner are both able to find end-to-end alignments for about ∼95% of the reads. STAR and PuffAligner are the fastest tools, with STAR being somewhat faster than PuffAligner. On the other hand, PuffAligner is able to align more reads than STAR, while requiring less than half as much memory. The memory usage of Bowtie2 is the smallest, since Bowtie2’s index does not contain a hash table. However, this comes at the cost of having the longest running time compared to other methods. Overall, PuffAligner benefits from the fast query of hash based indices while its run-time memory usage, which is mostly dominated by the size of the index, is significantly smaller than other hash based aligners. Although deBGA’s index is based on the de Bruijn graphs, similar to the Pufferfish index, the particular encoding for it is not as space-efficient as that of Pufferfish.

To look more closely how the mappings between the tools differ, we investigate the agreement of the reads which are mapped by each tool and visualize the results in an upset plot in fig. 1 using the UpsetR library^24^. We are only comparing the three methods which perform end-to-end alignment in this plot, since outliers from the local alignments computed by deBGA would otherwise dominate the plot. The first bar shows that the majority of the reads are mapped by all three tools.The next largest set represents the reads which are only mapped by Bowtie2 and PuffAligner. All the other sets are much smaller compared to the first two sets. This fact illustrates that the highest agreement in the aligners is between Bowtie2 and PuffAligner. Exploring a series of individual reads from the smaller sets in the upset plot, suggests that some of these differences happen as a result of small differences in the scoring configuration, while some result from different search hueristics adopted by the different tools. Supplementary fig. S1 shows the coherence between the alignments reported by the tools by also including the exact location to which the reads are aligned in the reference.

**Figure 1:**
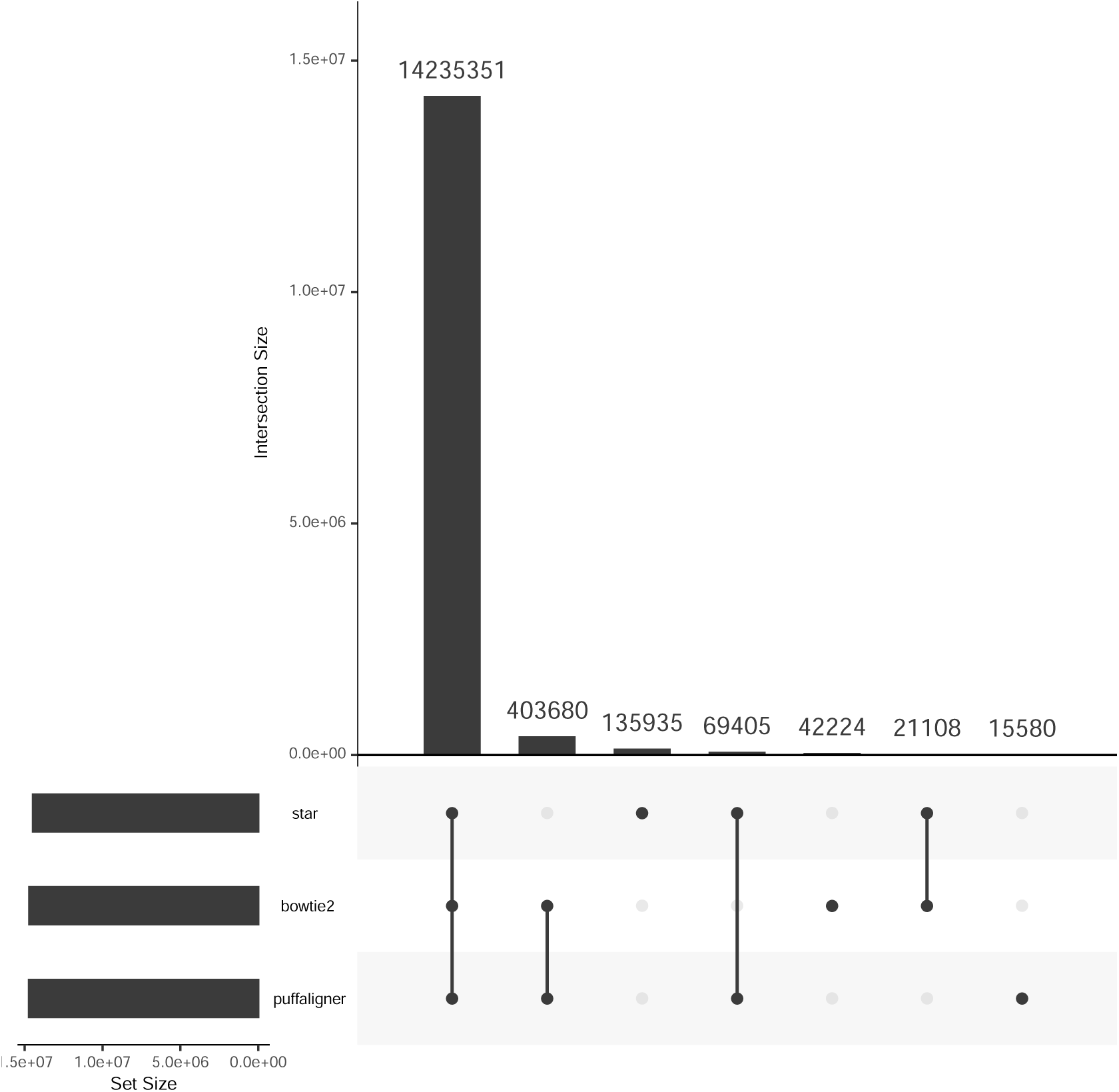
Upset plot showing the agreement of the alignments found by different tools

### 2.3 Alignment of simulated DNA-seq reads in the presence of variation

To further investigate the accuracy of the aligners, we used simulated DNA-seq reads.One of the main differences between simulated reads and experimental reads is that simulated reads are often generated from the same reference sequences to which they are aligned, with the only differences being due to (simulated) sequencing error. While (simulated) sequencing error prevents most reads from being exact substrings of the reference, it actually does not tend to complicate alignment too much. On the other hand, while dealing with experimental data, the genome of the individual from which the sample is sequenced might include different types of variations with respect to the reference genome to which we are aligning^25^. Therefore, it is desirable to introduce variations in the simulated samples, and to measure the robustness and performance of the different aligners in the presence of the variation. Mason^26^ is able to introduce different kinds of variations to the reference genome, such as SNVs, small gaps, and also structural variants (SV) such as large indels, inversions, translocations and duplications. We use Mason to simulate 9 DNA-seq samples with different variation rates ranging from 1*e* − 7 to 1*e* − 3. Each sample includes 1*M* paired-end Illumina reads of 100bp length from chromosome 21 of the human genome, ensembl release 98^‡^.

For this analysis, we do not restrict the aligners to only report concordant alignments, since the structural variations in the samples can lead to valid discordant alignments, such as those on the same strand or with inter-mate distances larger than the maximum fragment length. To be specific, we do not use the options which limit Bowtie2 and PuffAligner to report only concordant alignments, in addition, we use the option “`--dovetail`” in Bowtie2 to consider dovetail pairs as concordant pairs.

The alignments reported by deBGA already include discordant pairs and also orphan mappings. Furthermore, To remove any restrictions on the fragment length in the alignments reported by deBGA, we set the minimum and maximum insert size, respectively to 0 and the 50000, since setting a larger value resulted in the tool running into segmentation fault.

To allow dovetail pairs and also larger gaps between the pairs in STAR, we use the following options: “`--alignEndsProtrude 1000000 ConcordantPair`”, “`--alignMatesGapMax 1000000`”. By default there is not a specific option in STAR for allowing orphan alignment of paired end reads. Instead, we can increase the number of allowed mismatches to be as large as one end of the read by using the following options: “`--outFilterMismatchNoverReadLmax 0.5`”, “`--outFilterMismatchNoverLmax 0.99`”, “`--outFilterScoreMinOverLread 0`”, “`--outFilterMatchNminOverLread 0`”.

For each sample, Mason produces a SAM file which includes the alignment of the simulated reads to the original, non-variant version of the reference — the version which was used for building the aligner’s indices in this experiment. Based on the alignments reported in the truth file, some reads did not have a valid alignment to the original reference. This was the result of a high rate of variations at some sequencing sites. We called the set of reads that, according to the truth SAM file, were aligned to the original reference as compatible reads.

We compared the performance of aligners based upon how well they are able to align the compatible reads. We computed the precision and recall of the alignments reported for these reads as follows. True positives are considered the reads that are mapped by the aligner to the same location stated by the truth file. Then, recall is computed by dividing the number of true positives by the number of all compatible reads. Furthermore, we considered an alignment as a false positive in two different cases. First, an alignment was considered discordant if the reported alignment had a large edit distance (larger than 25) for the non-compatible reads. Second, in the case that an aligner reported an alignment to a location other than the one in the truth file, it was considered as a false positive if the edit distance of the reported alignment is greater than the edit distance of the true alignment. Having defined the set of TP and FP for the alignments, and also having considered the set of all compatible reads as the set we are trying to recover, we computed precision and recall for the set of alignments reported by each aligner.

Figure 2 shows the precision and recall of the aligners for different samples. According to fig. 2, for lower variation ratios up until 10*e* − 5, most of the tools are able to make accurate alignment calls with a high specificity. As the variation ratio introduced in the sample is increased, all the tools start to have lower precision and recall. deBGA and STAR perform worse in higher variation samples, as they fail to recover the true alignment for more reads, while Bowtie2 and PuffAligner are able to align most of the reads to their true location on the original reference.

**Figure 2:**
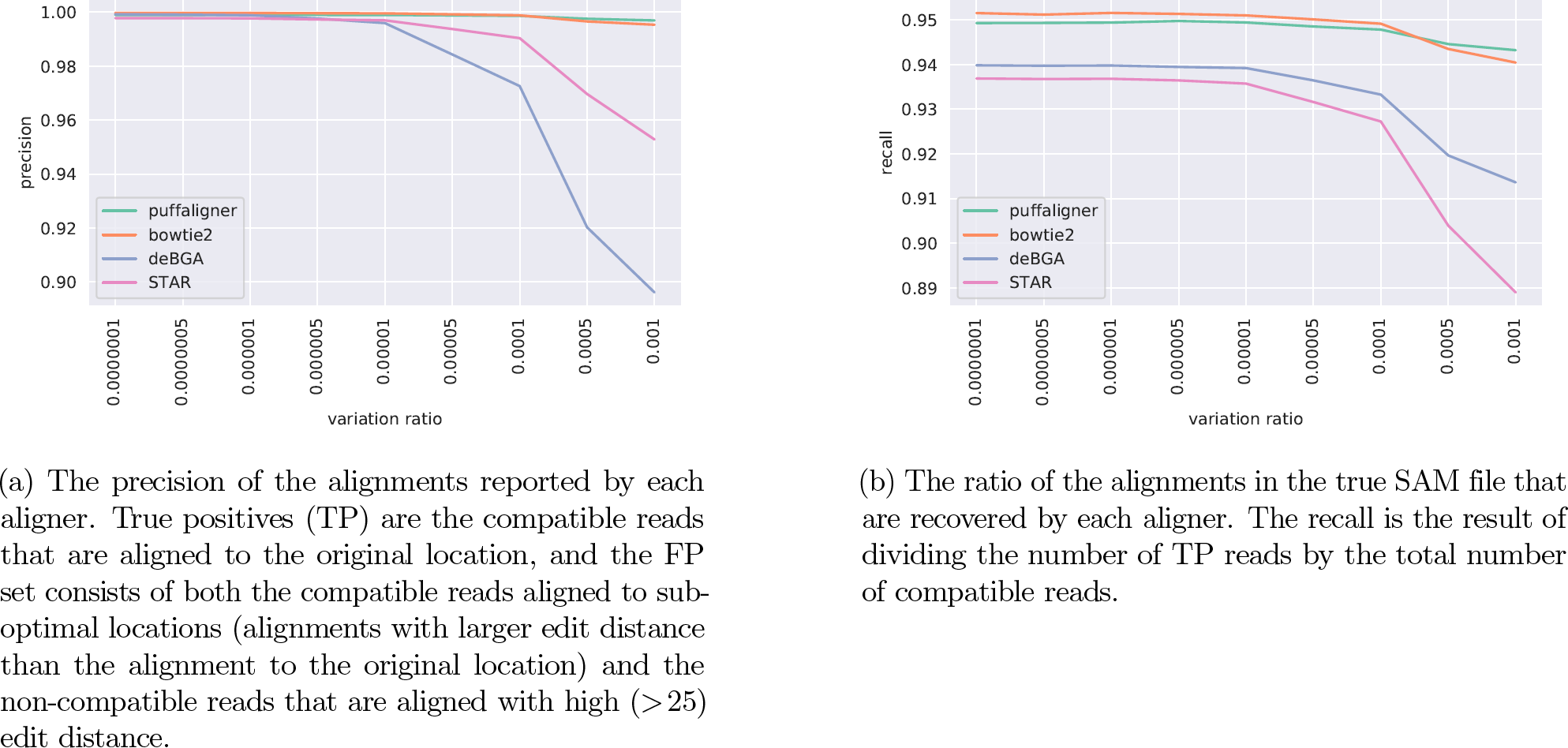
Comparing the accuracy of aligners in the presence of different rates of variations in the reference genome

These results show that PuffAligner’ accuracy is stable in the face of variation which makes the tool suitable for datasets that are known to have substantial variation, such as when aligning reads to microbial genomes where the specific sequenced strain may not be represented in the reference set.

### 2.4 Quantification of transcript abundance from RNA-seq reads

Mapping sequencing reads to target transcriptomes is the initial step in many pipelines for reference-based transcript abundance estimation. While lightweight mapping approaches^12,27^ greatly speed-up abundance estimation by, in part, eliding the computation of full alignment between reads and transcripts, there is evidence that alignments still yield the most accurate abundance estimates by providing increased sensitivity and avoiding spurious mappings^25,28,29^. Thus, the continued development of efficient methods for producing accurate transcriptome alignments of RNA-seq reads remains a topic of interest. In this section, we compare the effect of alignments produced by each tool on the accuracy of RNA-seq abundance estimation.

We generated 9,968,245 paired-end RNA-seq reads using the polyester^30^ read simulator. The reads are generated by the `simulate experiment countmat` module in polyester. The input count matrix is calculated based on the estimates from the Bowtie2-Salmon pipeline on the sample `SRR1085674` (where reads are first aligned with Bowtie2 and then the alignments are quantified using Salmon). This sample is a collection of paired-end RNA-seq reads sequenced from human transcriptome using an Illumina HiSeq^31^. The human transcriptome from gencode release (33) is used to build all the aligners’ indices. Also, for building STAR’s index in the genome mode, the human genome and the comprehensive gene annotation (main annotation file) is obtained from the same release of gencode.

As the reads in this experiment are RNA-seq reads sequenced from the human transcriptome, it is important to account for multi-mapping, as often, a read might map to multiple transcripts which share the same exon or exon junction. This property makes the direct evaluation of performance at the level of alignments difficult. Therefore, a typical approach in evaluating the accuracy of the transcriptomic alignments is to assess the accuracy of downstream analysis such as abundance estimations by computing the correlation and relative differences of the estimates with the true abundance of the transcripts. To compare the accuracy of each tool we give the alignments produced by each aligner, which are in the SAM format, as input to Salmon to estimate the transcript expressions.

PuffAligner, by default, outputs up to 200 alignments with an alignment score greater than 0.65 times the best alignment score, i.e., the alignment for the read in the case that all bases are perfectly matched to the reference. To enable the multi-mapping to take into account the characteristics of alignment to the transcriptome, Bowtie2 is run with the option `-k 200` which lets the tool output up to 200 alignments per read. The value of 200 is adopted from the suggested parameters for running RSEM^32^ with Bowtie2 alignments. We note that running Bowtie2 with this option makes the tool considerably slower than the default mode, as many more alignments will be computed and output to the SAM file under this configuration. For both Bowtie2 and PuffAligner, and also for STAR by default, orphan and discordant mappings are not allowed.

We ran STAR with the `‘ENCODE’` options, which are recommended in the STAR manual for RNA-seq reads. STAR is also run in two different modes, one is by building the STAR index on human genome, while it is also provided a GTF file for gene annotation. In this mode, STAR performs spliced alignment to the genome, then projects the alignments onto transcriptomic coordinates. The other mode is building the STAR index on the human transcriptome directly, which allows STAR to align the RNA-seq reads directly to the transcripts in an unspliced manner. We chose to run STAR in the transcriptomic mode as well, since we find that it yields higher accuracy, though this increases the running time of STAR.

The deBGA index is built on the transcriptome, as are the Bowtie2 and PuffAligner indices, since these tools do not support spliced read alignment. deBGA is run in the with options `-o 200 -x 200`, which nominally has the same effect as `-k 200` in Bowtie2, according to the documentation of deBGA.

Accuracy of abundance estimation by Salmon, when provided the SAM output generated by each aligner, is displayed in table 2. The timing and memory benchmarks provided in this table is only for the alignment step. Alignments produced by PuffAligner, Bowtie2 and STAR in the transcriptomic mode produce the best abundance estimates. deBGA’s output alignments are not suitable for any abundance estimation as many reads are aligned only to the same strand which are later filtered during the abundance estimation by Salmon, so we could not provide a meaningful correlations for abundance estimation using deBGA’s alignments. Aligning the reads by STAR to genome and then projecting to transcriptomic coordinates does not generate as high correlation as directly aligning the reads to the transcriptome by STAR. However, we note that, as described by Srivastava et al. ^25^, there are numerous reasons to consider alignment to the entire genome that are not necessarily reflected in simulated experiments. While the memory usage by PuffAligner is only 2 fold larger than memory used by Bowtie2, it computes the alignments much more quickly.

**Table 1:** The performance of different tools for aligning experimental DNA-seq reads. The time reports are benchmarked after warming up the system cache so that the influence of index loading time is mitigated.

| aligner | mapping-rate(%) | time (mm:ss) | memory (GB) |
| --- | --- | --- | --- |
| PuffAligner | 95.58 | 6:14 | 13.09 |
| deBGA | 99.75 | 10:46 | 41.04 |
| STAR | 93.88 | 4:29 | 30.36 |
| Bowtie2 | 95.44 | 16:15 | 3.50 |

**Table 2:** Abundance estimation of simulated RNA-seq reads, computed by Salmon, using different tools’ alignment outputs. The time and memory are only for the alignment step of each tool and the time for abundance estimation by Salmon is not considered.

| aligner | spearman | MARD | time (mm:ss) | memory (GB) |
| --- | --- | --- | --- | --- |
| PuffAligner | 0.92 | 0.05 | 1:17 | 2.54 |
| deBGA | N/A | N/A | 5:19 | 9.96 |
| STAR- transcriptome | 0.92 | 0.05 | 1:57 | 8.73 |
| STAR- genome | 0.90 | 0.06 | 3:30 | 32.57 |
| Bowtie2 | 0.92 | 0.05 | 32:59 | 1.15 |

According to the results in table 2 PuffAligner is the fastest aligner in these benchmarks, and the accuracy as high as Bowtie2 and STAR for aligning RNA-seq reads. Here, PuffAligner leads to the most accurate abundance estimates, while being 30 times faster than Bowtie2. Moreover, The memory usage is much less than other fast aligners such as STAR.

### 2.5 Alignment to a collection of microorganisms — simulated short reads

To demonstrate the performance and accuracy of PuffAligner for metagenomic samples, we designed two different experiments. One main property of metagenomic samples is the high similarity of the reference sequences against which one typically aligns, where a pair (or more) of references may be more than 90% identical. The first experiment we designed for this scenario, to specifically evaluate issues related to this challenge, we call the “single strain” experiment. Additionally, metagenomic samples also have the property of containing reads from a variety of genomes, some of which are not even assembled yet – and hence unknown. This leads to the second experiment, which we call the “bulk” experiment, that compares the aligners in the presence of a high variety of `species` in the sample in addition to the high similarity of references.

For simplicity and uniformity, all the experiments have been run in the concordant mode for both PuffAligner and Bowtie2 (both of which support such an option), disallowing orphans and discordant alignments. All aligners are run in three different confiurations, allowing three specific maximum numbers of alignments per fragment; 1 (primary output with highest score, breaking ties randomly), 20, and 200. PuffAligner and STAR, as the only tools that support this option, also are run in the *bestStrata* mode. In this mode, the aligner outputs all *equally-best* alignments for a read with highest score without the limitation on number of reported alignments. This option is inspired by the similarly-named option in Bowtie1^33^. However, unlike Bowtie1, PuffAligner and STAR only make a best-effort attempt to find the score of the best stratum alignments, and do not guarantee to find the best stratum (though the cases in which they fail to seem to be exceedingly rare). This option is especially useful in the metagenomic analyses, as we will report only the best-score alignments without having an arbitrary limitation on the number of allowed alignments. This allows proper handling of highly multi-mapping metagenomic reads. In other words, using this option, one can achieve a high sensitivity without the need to hurt specificity. The details of each experiment is explained in the following sections.

#### 2.5.1 A single-strain Experiment

For this experiment, we download the viral database from NCBI, and choose three similar coronavirus genomes. This set includes one of the recently-uploaded samples from Wuhan^34,35^. We select three very similar viral genomes to simulate reads from, which are: NC_045512.2, NC_004718.3, and NC_014470.1. There are also a lot of literature discussing the similarity in sequence and behavior for these three species of coronavirus^36^–38. The first is the complete genome for severe acute respiratory syndrome coronavirus 2 isolate Wuhan-Hu-1 known as Covid19 with length of 29,904 bases. NC_004718.3 is the ID of SARS coronavirus complete genome (length: 29,752) and finally, NC_014470.1 is a Bat coronavirus BM48-31/BGR/2008 complete genome (length: 29,277).

We use Mason^26^ to generate three simulated samples, each sample contains 500,000 reads only from one of the three viral references we mentioned earlier. Then, reads were aligned back to the database of viral sequences using each of the four aligners. The results are shown in table 3 for covid19 and table S4 for the other two simulations.

**Table 3:** Alignment Distribution for 500000 simulated reads from reference sequence NC_045512.2 (known as covid19). The best specificity is achieved by PuffAligner in `bestStrata` mode (as well as the primary mode). In this simulated sample, many alignments are not ambiguous, resulting in the good performance observed when using only primary alignments. However, typically in metagenomic analysis, many equally-good alignments exist, and selecting only one is equivalent to making a random choice.

| Alignment Mode | Tool | NC_045512.2<br>(Covid19) | NC_004718.3<br>(SARS Coronavirus) | NC_014470.1<br>(Bat Coronavirus) | Other Assembled<br>References |
| --- | --- | --- | --- | --- | --- |
| Primary | PuffAligner | 500,000 | 0 | 0 | 0 |
|  | Bowtie2 | 499,981 | 18 | 0 | 0 |
|  | STAR | 499,999 | 0 | 0 | 0 |
|  | deBGA | 499,991 | 0 | 0 | 9 |
| Up to 20 | PuffAligner | 500,000 | 134 | 46 | 0 |
|  | Bowtie2 | 499,999 | 21,461 | 2,311 | 0 |
|  | STAR | 499,999 | 0 | 0 | 0 |
|  | deBGA | 499,991 | 0 | 0 | 9 |
| Up to 200 | PuffAligner | 500,000 | 134 | 46 | 0 |
|  | Bowtie2 | 499,999 | 21,461 | 2,311 | 0 |
|  | STAR | 499,999 | 0 | 0 | 0 |
|  | deBGA | 499,991 | 0 | 0 | 9 |
| Best strata | PuffAligner | 500,000 | 0 | 0 | 0 |
|  | STAR | 499,999 | 0 | 0 | 0 |

As the results show, the alignments of all aligners, except for deBGA, are distributed only across the three references of interest out of all the reference sequences in the complete viral database. deBGA reports only a few alignments to a forth virus. In general, all of the aligners do a good job of reporting the correct alignment among the returned alignments for each read. Here, we are more interested in exploring how sub-optimal alignments are computed and filtered under different settings when aligning to a collection of very similar genomes. The results show that all tools have very high sensitivity even when considering only a single (primary) alignment per read. As we allow more alignments to be reported, the sensitivity increases and quickly levels off for all the tools. On the other hand, more alignments are generated and Bowtie2, in particular, generates a considerable number of extra alignments as the maximum number of allowed reported alignments is increased. However, the results do not change when allowing more than 20 alignments, which means no more than 20 alignments ever pass the alignment score threshold for these reads in the viral database for any of the tools we are testing.

The results indicate that, when allowing more than one alignment to be reported for every read, Bowtie2 tends to report a large number of sub-optimal (yet, still valid) alignments compared to other tools. These are alignments that are accepted within the alignment score threshold, but are to another target than the one from which the read truly originates. Generating these sub-optimal alignments is in no way wrong, but it has a non-trivial computational cost, as shown in figure 3a, even if these alignments are not used in downstream analysis. Further, the score of the best alignment for each read is specific to that read and not known ahead of time, meaning that this situation cannot be completely addressed simply by setting more stringent parameters for which alignment scores should be allowed. This behavior of Bowtie2 gives the other tools a computational advantage when the user only truly requires the set of equally-best alignments for each read.

**Figure 3:**
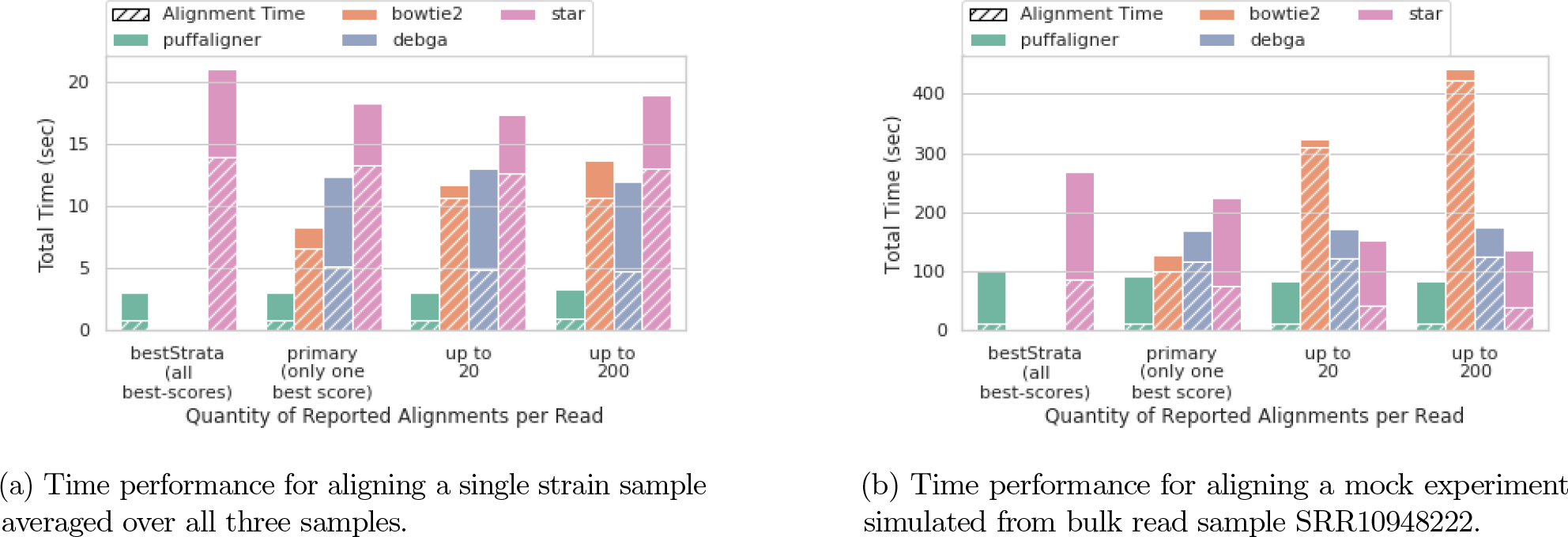
Time performance of different aligners on the two microbiome experiments. In 3a, the results are averaged over the three alignment processes for the samples covid19, sars, and bat200, each having ∼1*M* paired-end reads. In 3b the performance shown is for aligning reads in mock sample simulated from SRR10948222 with 5*M* paired-end reads. As shown in the bulk experiment, the alignment for Bowtie2 increases when asking for more alignments per read while the other tools show a constant alignment time scaling over number of reads. The dashed area shows fraction of the time spent purely on aligning reads where the remaining portion is the time required for index loading. PuffAligner is the fastest tool in this experiment, yet most of its time is still dedicated to loading the index.

Interestingly, there is one read that all tools, except for PuffAligner fail to properly align. Inspecting this alignment reveals it is a valid alignment within the range of the acceptable scoring threshold, and it is unclear why it is not discovered by the other tools. Overall, the aligners tested perform very well here in reporting the true strain of origin without reporting too many extra alignments. Interestingly, despite changing the parameters to allow more alignments, STAR tends to return the same set of alignments under all configurations in this experiment. Figure 3 shows that PuffAligner has the lowest running time, even when the number of allowed alignments per read increases.

#### BestStrata Mode

In this small example, all tools showed good sensitivity (and PuffAligner and STAR showed near-perfect sensitivity) even when reporting only a single-alignment per read. This experiment is, of course, an atypically small test for multi-mapping read. In in larger samples, with reads deriving from more organisms and a larger database of references, permitting more alignments usually yields non-trivial improvements in sensitivity. To control the rate of reporting sub-optimal alignments, PuffAligner supports the “best strata” option – also available to STAR, which allows only the alignments with the best calculated score to be reported (as a replacement for maximum allowed number of alignments). Using this option, PuffAligner achieves full specificity and sensitivity in this experiment 3. The same results are achieved for the other two simulated single-strain samples shown in the supplementary table S4. We further demonstrate the positive impact of this option on the alignment of bulk metagenomic samples in the next section.

### 2.5.2 Experiments with a mixture of organisms

We chose a random set of 4000 complete bacterial genomes downloaded from the NCBI microbial database and constructed the indices of PuffAligner, Bowtie2, STAR, and deBGA on the selected genomes. Supplementary Table S2 shows the time and memory required for constructing each of the indices, in addition to the size of the final index on disk. Overall, PuffAligner and Bowtie2 show a similar trend in time and memory requirements, while STAR and deBGA require an order of magnitude more memory. In terms of the final index size, Bowtie2 has the smallest index, PuffAligner has the second-smallest, and STAR has the largets.

For simulating a bulk metagenomic sample, we generated a list of simulated whole genome sequencing (WGS) reads through the following steps:

- Select a real metagenomic WGS read sample
- Align the reads of the chosen real experiment to the 4000 genomes using Bowtie2, limiting Bowtie2 to output one alignment per read.
- Choose all the references with count greater than *C* from the quantification results. This defines the read distribution profile that we will use to simulate data.
- For each of the expressed references, use Mason^26^, a whole genome sequence simulator, to simulate 100*bp* paired-end reads with counts proportional to the reported abundance estimates so that total number of reads is greater than a specified value *n*. In this step we ran Mason with default options.
- Mix and shuffle all of the simulated reads from each reference into one sample which is used as the mock metagenomic sample.

We selected three Illumina WGS samples that are publicly available on NCBI. A soil experiment with accession ID `SRR10948222`^39^ from a project for finding sub-biocrust soil microbial communities in the Mojave Desert. The sample has ∼27*M* paired-end reads, containing a mixture of genomes from various genera and families. However, less than 200*k* of the reads in the sample were aligned to the strains present in our database, leading the selection of 98 species from a variety of genera. We scaled the read counts in the simulation to ∼50*M* reads. The other two selected samples are `SRR11283975` and `SRR11496426` the details of which are explained in supplementary S3. In this section we only report the performance of the tools on the first sample. The analysis results for the other samples (which shows similar relative accuracy and performance for different tools) are provided in S5.

The assessment of “accuracy” directly from the aligned reads is not a trivial task. Due to the high rate of multi-mapping in these simulated samples, and due to the fact that multiple references can produce alignments of the same quality as the “true” origin of the read, we calculate the accuracy by comparing the true and estimated abundances using a quantification tool (in this case, Salmon) rather than by comparing the read alignments directly. In Table 4 the accuracy metrics are calculated over the abundance estimations obtained using the alignments produced by running the aligners in the different modes specified. The list of metrics for metagenomic expression evaluations have been chosen to be similar to previous work such as in Bracken^40^ and Karp^41^.

**Table 4:** Accuracy of abundance estimation with Salmon using alignments reported by each aligner for the mock sample simulated from a real sample with accession ID SRR10948222. We have run all the aligners in three main modes; allowing only one best alignment with ties broken randomly (Primary), up to 20 alignments reported per read, and up to 200 alignments reported per read. PuffAligner and STAR support a fourth mode that allows reporting all equally best alignments (bestStrata). This option improves the performance while maintaining, or even slightly improving, the accuracy of the results.

| Alignment Mode | Tool | Spearman | MARD | MAE | MSLE |
| --- | --- | --- | --- | --- | --- |
| Primary | PuffAligner | 0.69 | 0.028 | 1.39 | 0.08 |
|  | Bowtie2 | 0.58 | 0.053 | 2.91 | 0.15 |
|  | STAR | 0.727 | 0.023 | 1.493 | 0.05 |
|  | deBGA | 0.28 | 0.616 | 656.08 | 6.53 |
| Up to 20 | PuffAligner | 0.9 | 0.006 | 0.40 | 0.01 |
|  | Bowtie2 | 0.85 | 0.01 | 0.22 | 0.01 |
|  | STAR | 0.929 | 0.004 | 0.303 | 0.00 |
|  | deBGA | 0.28 | 0.573 | 637.60 | 5.65 |
| Up to 200 | PuffAligner | 0.97 | 0.002 | 0.36 | 0.00 |
|  | Bowtie2 | 0.99 | 0.001 | 0.19 | 0.00 |
|  | STAR | 0.929 | 0.004 | 0.299 | 0.00 |
|  | deBGA | 0.28 | 0.571 | 637.83 | 5.55 |
| Best strata | PuffAligner | 0.97 | 0.002 | 0.36 | 0.00 |
|  | STAR | 0.929 | 0.004 | 0.3 | 0.00 |

The metrics selected are *Spearman Correlation, Mean Absolute Relative Difference (MARD), Mean Absolute Error (MAE)*, and *Mean Squared Log Error (MSLE)*. Each metric measures different characteristics of the predicted versus true abundance estimates. For example, lower MARD indicates better distribution of the reads among the references relative to the abundance of each reference, while MAE shows the quality of the distribution of the reads in a more absolute way regardless of the difference between the abundance of the references. In this case, one misclassified read has the same impact on the MAE metric both for a high-abundance and low-abundance reference. The definition of each of these metrics is provided in equation 2 in the supplementary material.

This experiment leads to three main observations. First, regardless of the alignment mode, quantifications derived from the deBGA alignments seem to lead to systematic underestimation of abundance. However, PuffAligner, STAR and Bowtie2, show very similar behavior with respect to accuracy. STAR is the best in primary mode as well as when allowing 20 alignments, closely followed by PuffAligner. When allowing up to 200 alignments per read, Bowtie2 tends to yield the most accurate abundances, again with PuffAligner being the close runner-up. These results demonstrate that PuffAligner is a reliable alignment tool showing a stable pattern of being comparable to the best aligner under all the scenarios tested. That is, the good performance of PuffAligner is robust across a variety of different parameter settings.

Moreover, due to the nature of the metagenomic data — the high degree of ambiguity and multi-mapping — we expect to see improvement in the accuracy metrics as more alignments are reported per read, as this leads to a higher recall. While STAR’s accuracy changes only slightly from 20 alignments to 200 alignments (only improving MAE) the results for PuffAligner and Bowtie2 improve considerably when allowing more alignments per read. However, this higher accuracy comes in the cost of alignment time for Bowtie2. As shown in figure 3, section 3b, Bowtie2 alignment time increases sharply when allowing more alignments per read, while PuffAligner exhibits only small changes in alignment time regardless of the maximum number of alignments being reported per read. The difference becomes especially evident when allowing up to 200 alignments per read, where PuffAligner is 4 times faster than Bowtie2. Additionally, in experimental data, many of the alignments reported do not necessarily have high quality, and only appear in the output as one of the 200 alignments for the read. In fact, we note the similar accuracy achieved by PuffAligner in *bestStrata* mode compared to when we allow up to 200 alignments per read. This observation is also consistent across the other two simulated samples in the supplementary table S5, in those cases with PuffAligner being the most accurate aligner in different modes for both samples.

Overall, these results along with other similar experiments in supplementary table S5 indicate that PuffAligner is a sensitive and fast aligner. Specifically PuffAligner exhibits similar accuracy (and is sometimes more accurate) as well-known aligners like Bowtie2 and STAR. On these data, it exhibits memory requirements close to those of the memory-frugal Bowtie2, while being much faster.

### 2.6 Scalability

Figure 4 represents how the construction time and index size of each tool scales over different types of sequences. The trend shows the effect of database size as well as redundancy and sequence similarity on the scalability of each of the tools. Tools such as PuffAligner and deBGA, which build a de Bruijn graph based index on the input sequence, specifically compress similar sequences into unitigs and therefore scale well for databases with high redundancy such as microbiomes. It is worth mentioning that Bowtie2 requires a switch from a 32-bit index to a 64-index as the total count of the input bases increases, which is another reason why the size is growing super-linearly.

**Figure 4:**
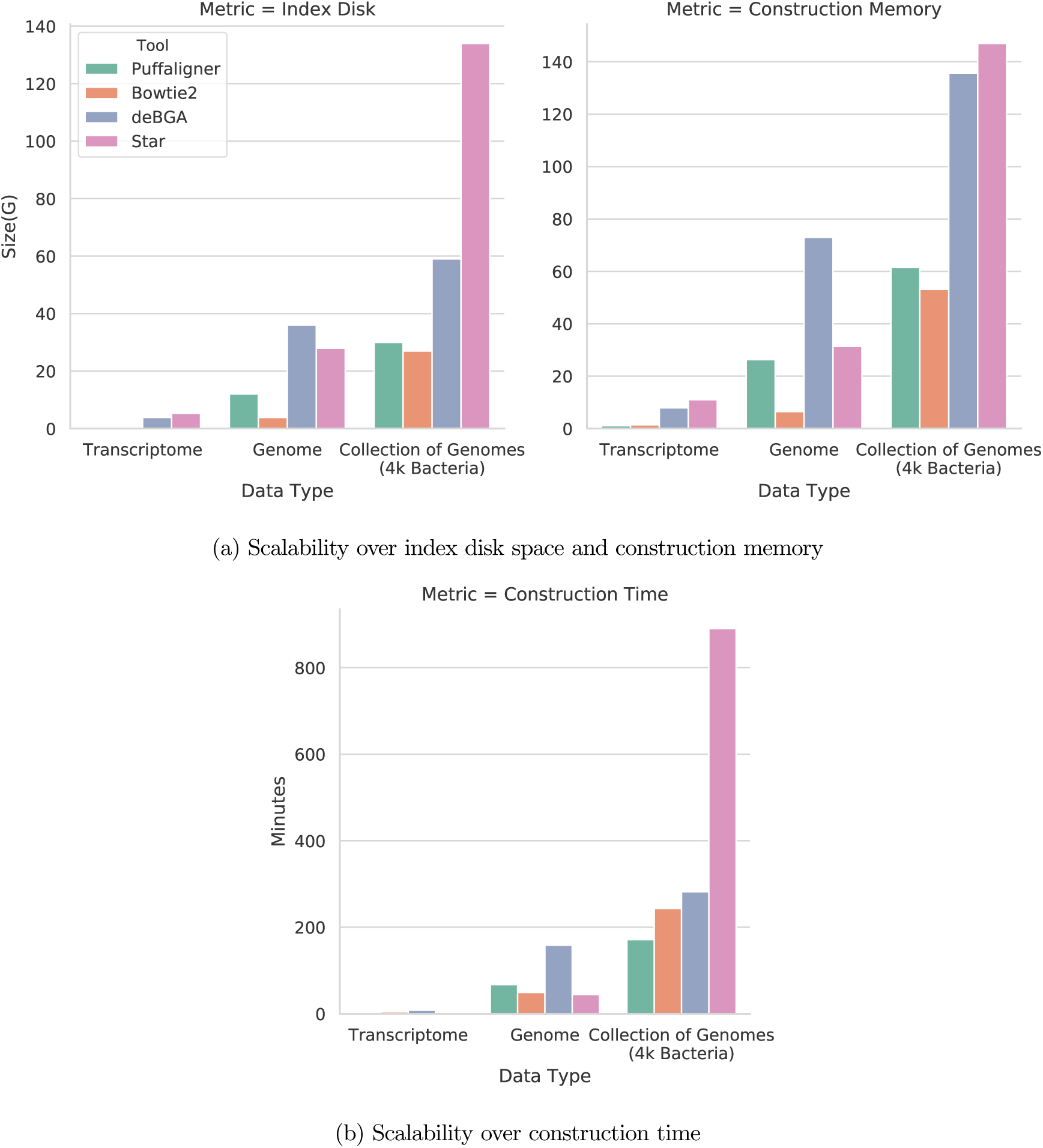
Scalability of different tools over the final index disk space, construction memory, and construction running time for three different datasets, human transcriptome (gencode version 33), human genome (GRCh38 primary assembly), and collection of genomes (4000 random bacterial complete genomes). All tools are run with 16 threads.

### 2.7 Comparing alignment to a light-weight pipeline like Kraken2 + Bracken

Here, we compare a pipeline for metagenomic abundance estimation comprised of aligning reads using PuffAligner followed by quantification with Salmon, to a *k*-mer -based abundance estimation method that consists of classifying reads with Kraken2 and estimating the abundances with Bracken. To perform this comparison, we construct an index over all bacteria, viral, archaea, and fungi obtained from NCBI taxonomy database^42^ on May 21, 2020 using both Pufferfish and Kraken2. We ran both approaches over 34 samples from different categories of metagenomic analyses. Ten of the samples are selected from non-human projects such as the “metasub project”^43^ as well as from submarine or soil samples^44^. The remaining samples are selected from the human metagenome project (hmp)^45^. The human samples are chosen from different tissue categories of plasma, tongue dorsal, gingiva, vaginal, and fecal. We then compare the abundance of the reported references under each of the two pipelines.

We run PuffAligner in two modes; the default mode, which only reports the best concordant alignments, and a less restrictive mode which will be explained. The default mode does not allow any orphan, discordant, or dovetail alignments, filters any mapping with alignment score less than 0.65 fraction of the maximum possible alignment score, i.e., when the read matches the reference perfectly, and for each read reports only alignments with the highest score (bestStrata). We also ran PuffAligner allowing orphans, discordant, and dovetail alignments, but keeping the rest of the parameters as default. We use Salmon to estimate the abundance of references from the alignments. While this is a reasonable pipeline for metagenomic abundance estimation, it is likely that the accuracy could be further improved by incorporating specific features of the metagenomic data in the abundance estimation step, such as the topology of the taxonomy tree and the expression of certain marker genes.

We also run Kraken2 in two different modes. First, we run it in the “default” mode, which allows all reads having any *k*-mer match to be classified. Second, we run it setting the confidence option to a value *c*, which prevents reporting reads that have a confidence lower than *c*. As per the authors’ definition^§^, this number is calculated based on the ratio of unique *k*-mers mapped to a taxa in the taxonomy tree over all the non-ambiguous *k*-mers of the read (*k*-mers not containing an `N` character). There is not a one-to-one correspondence between the confidence threshold here and the “minScoreFraction” in PuffAligner. However, we believe both of these options are necessary for providing more reliable abundance estimates, by removing the reads that exhibit poor evidence of deriving from an indexed references. Filtering orphaned, discordant, or dovetail reads is a feature only available to the alignment-based procedures and not *k*-mer counting approaches, since they do not jointly consider such properties of the reads containing the *k*-mers. That is why we provide both modes for PuffAligner, allowing and disallowing these sort of reads, to provide a fair comparison of the two methods.

In the absence of technical errors and variation from the reference genomes, if a sequencing read comes from a subset of references in the index, then there will be at least one exact match for it. However, due to the presence of both sequencing errors and biological variation of the organisms in the samples with respect to the reference genomes, a perfect alignment does not exist for most reads. Therefore, alignment-based methods try to find the sub-sequences in the reference which matches the read with a high alignment score (including both mismatches and gaps) and report these loci as the potential origin of the read. On the other hand, *k*-mer -based approaches break the read into its constituent *k*-mers and treat exact *k*-mer matches as evidence of a read originating from a reference. In most cases, there is a large degree of concordance between this approach and the alignment-based approaches for finding potential loci (i.e. the number of *k*-mer matches correlates well with alignment score). Thus, in general, such approaches result in highly-correlated abundance estimates with those produced by alignment-based approaches, and they also tend to be very fast and memory efficient. Nonetheless, such approaches still face challenges as the degree of sequence-similarity between strains (and species) can be very large and, simultaneously, the organisms present in a sample can show considerable sequence divergence from the strains in the reference database. As a result, *k*-mer -based approaches, while very sensitive, tend to sacrifices specificity in this type of data. In most cases, when no filter is applied to the taxonomic assignments, a *k*-mer -based approach overestimates the number of reads deriving from the reference strains in each sample.

To see how the reference abundance estimations compare across the two pipelines of Kraken2 and PuffAligner we look at the top 5 most highly-abundant (predicted) species for each sample and their abundance in supplementary figure S2. We run Kraken2 both with no confidence filter and filtering with a confidence value of .65, and compare the results with default PuffAligner (with minScoreFraction=.65). In certain samples, observe similar highly-abundant species discovered by Kraken2 (with no confidence filter) and PuffAligner that are reported as less abundant when processed with Kraken2 using a confidence threshold of 0.65 (e.g. Streptococcus gordonii for one of the subway samples). However, for other samples, applying the confidence threshold to the Kraken2 results brings the predictions closer to those of PuffAligner (e.g. Lactobocillus Crispatus for two vaginal sample). We do the same analyses at the genus level in figure S3 and there we observe similar inconsistencies.

To further investigate how the filtering of low quality reads in the two approaches compare, we look at the trend of the number of reads filtered in Kraken2 while applying different confidence thresholds with respect to the default PuffAligner in figure 5. We run Kraken2 with confidence values of .5, .65, .8, as well as .05 which is recommended by the authors for filtering low quality alignments for general purposes^¶^. We compare the results of running the Kraken2 + Bracken pipeline with PuffAligner + Salmon pipeline under both modes of PuffAligner (default and less restrictive) so that the effect of the read-pair-based filters (those taking into account read orphan and dovetail status) can be evaluated. The plots in the top row of figure 5 show the read count difference of default PuffAligner and Kraken2 applying different confidence values. The total read count varies widely across samples. Therefore, to have a better understanding of the results, we normalize the read count differences based on the PuffAligner reported read count in the two bottom plots of figure 5.

**Figure 5:**
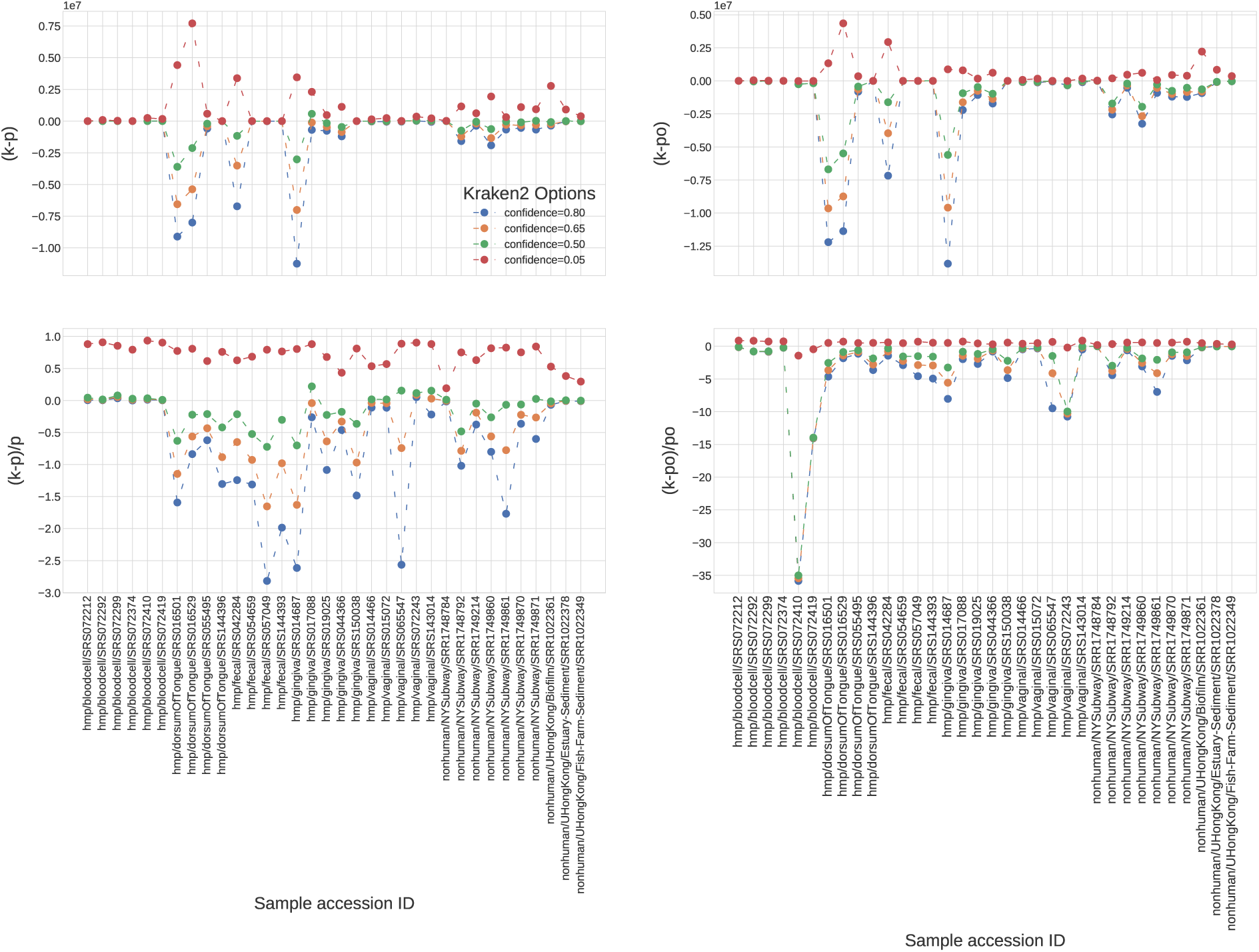
This plot shows the effect of applying different confidence values on the results of Kraken2 + Bracken pipeline relative to PuffAligner in two modes; default, and less restrictive mode (allowing orphan, discordant, or dovetail alignments). The abbreviations on the y-axis labels are defined as follows: k=Kraken2, p=PuffAligner (default), po=PuffAligner (allowing discordant alignments). Applying different confidence values to the Kraken2 pipeline results in an inconsistent pattern of assigned reads across samples when comparing to the PuffAligner read count (for both modes of PuffAligner (default on left and restrictive on right)). The count of filtered reads for both PuffAligner and Kraken2 depends on the sample and quality of the reads (the two plots on the top) as well as the filtered read counts relative to PuffAligner’s (the bottom plots). However, there is an observed inconsistency in how changing the confidence threshold of Kraken2 affects the number of filtered reads when comparing to the alignment-based approach of PuffAligner. In fact, not only is there no fixed confidence value that corresponds to the analogous alignment options, but further there is no corresponding value that provides a smooth consistent effect on the number of filtered reads. There are large changes from allocating many more read to allocating many fewer reads compared to PuffAligner in some samples, whereas the allocated read count barely changes in others.

As is observed, applying the confidence threshold may not cause a uniformly similar impact on the absolute read count for all samples. We expect to see a similar relative effect when applying the threshold over all the samples, i.e. the read count consistently becoming closer to or further from that obtained by the alignment-based approach. Although this is the observed trend for small confidence values which are less effective, the plots in figure 5 do not show a consistent pattern for larger confidence values in all the samples, where the application of the threshold in Kraken2 makes the reported read counts sometimes closer and sometimes farther from the alignment-based pipeline. As might be expected, the read counts reported in Kraken2 are further from the PuffAligner pipeline in the default mode versus when allowing orphans and discordant alignments in the less restrictive mode (as can be seen when comparing the two columns of figure 5).

To summarize, through the experiments in this section, we observe that the *k*-mer -based approaches sometimes result in different reported highly-abundant references than alignment-based approach and, crucially, applying a score filter to the *k*-mer -based approaches does not always make the abundance predictions for all species closer to the alignment-based predictions. The efficacy of the score filter in the *k*-mer -based approaches may depend on the technical and biological details particular to that sample, so that the score filter that best matches the alignment-based approach in one sample may not be the same score filter that best matches the alignment-based approach in other samples. This is, perhaps, expected, as the alignment-based approach is taking into account more information, both within and across the ends of a paired-end fragment, than simply the number of matching *k*-mers.

In this sense, we believe that a sensitive and efficient aligner like PuffAligner, paired with a capable abundance estimation tool, provides a pipeline for metagenomic abundance estimation that, while sacrificing some speed, is likely to be more accurate and robust than *k*-mer -based approaches.

## 3 Discussion & Conclusion

In this paper we introduce PuffAligner, an aligner suitable for the contiguous alignment of short-read sequencing data. We demonstrate its use in aligning DNA-seq reads to the genome of a single species, aligning RNA-seq reads to the transcriptome, and aligning DNA-seq reads from metagenomic samples to a large collection of references. It is built on top of the Pufferfish index, which constructs a colored compacted de Bruijn graph using the input reference sequences. PuffAligner begins read alignment by collecting unique maximal exact matches, querying *k*-mers from the read in the Pufferfish index. The aligner then chains together the collected uni-MEMs using a dynamic programming approach, choosing the chains with the highest coverage as potential alignment positions for the reads. Finally, PuffAligner is able to efficiently compute alignment, exploiting information from long matches in the chains and making use of an alignment cache to avoid redundant work.

We compared the accuracy and efficiency of PuffAligner against two widely-used alignment tools, Bowtie2 and STAR, that perform unspliced and (optionally) spliced alignments of reads, respectively. We also compare the results against deBGA, an aligner that also utilizes an index built over the compacted de Bruijn graph.

We analyze the performance of these tools on both simulated and experimental DNA and RNA sequencing datasets. The accuracy of PuffAligner is comparable to Bowtie2, which exhibits very high alignment. PuffAligner generally performs better than STAR and deBGA (though, unlike STAR, none of these other tools currently support spliced read alignment). In terms of speed and memory, PuffAligner reaches a tradeoff between the relatively high memory usage of STAR and deBGA and the slower speed of Bowtie2. Hence, while the memory requirement of PuffAligner is more than that of Bowtie2, the speed gain is significant. In the tests performed in this manuscript, PuffAligner is almost always the fastest tool (with the exception being that STAR is faster when aligning unspliced DNA-seq reads to a single human genome).

An additional advantage of the Pufferfish index utilized in PuffAligner is that it can be built on a mixed collection of genomes, transcriptomes, or both. This feature is already utilized in a specific pipeline for RNA-seq quantification that makes use of a joint index over the genome and transcriptome^25^. The analysis shows that specificity of alignments in such a case can be improved by filtering from quantification reads that are better aligned to some genomic locus that is not present in the transcriptome.

Furthermore, the nature of the Pufferfish index, that explicitly factorizes out highly-repetive sequence, coupled with the fast (and repetition-aware) alignment procedure of PuffAligner makes it a particularly useful for indexing and aligning to a highly similar collection of sequences. This potentially makes it a good match for metagenomic analyses.

We have provided a proof of concept for such a PuffAligner-based metagenomic analysis pipeline, and plan to build a more sophisticated and fully-featured metagenomic analysis framework around PuffAligner in the future.

## 4 Materials and Methods

PuffAligner is an aligner built on top of the Pufferfish indexing data structure. Pufferfish is a space-efficient and fast index for the colored compacted de Bruijn graph (ccdBg). A colored compacted de Bruijn graph is a graph whose vertices (strings) are the compacted non-branching paths of the underlying de Bruijn graph, with the restriction that each node also have the same color set (set of reference sequences in which it appears). The nodes in the colored compacted de Bruijn graph are referred to as unitigs. Each unitig can be mapped to a list of *<*reference ID, position, orientation≥ tuples that describe exactly how this subsequence appears in the unlderying collection of references. The basic query operation in the Pufferfish index is to query a *k*-mer from the input sequence against the index. Given this query, the pufferfish index returns the unique position (and orientation) where this *k*-mer appears in the colored compacted de Bruijn graph (or a sentinel value if this *k*-mer does not occur). This match between the query and the graph can then be easily “unpacked” into the implied list of matches with the underlying references by finding all of the places that the matched unitig appears in the reference sequences and translating the relative position within the unitig into the corresponding reference position (and adjusting the orientation if necessary). The output of this step is then a list of all of the reference sequences, positions, and orientations where this exact match occurs. While *k*-mer query is the basic operation performed by the index, we actually do not use *k*-mer matches directly, and instead extend the initial match into unique maximal exact matches (uni-MEMs).

Specficially, each *k*-mer match is extended simultaneously in both the query and reference to obtain a longer exact match. The exact matches to the unitigs, called uni-MEMs, are then projected to the positions on the references associated to that unitig. Then, uni-MEMs are aggregated into MEMs (described below) on each reference, and the chains of MEMs with the highest score are selected. In the case of paired-end reads, the chains of the left and right ends are paired with respect to their distance, orientation, etc. Finally, rather than fully aligning each query sequence to the anchored position on the reference, only the sub-sequences from the query that are not part of the uni-MEMs (exact matches) are aligned to the reference; we call this procedure the between-MEM alignment. Each of these steps are explained in detail in the following sections.

### 4.1 Exact matching in the Pufferfish index

The pufferfish index provides PuffAligner with an efficient index for *k*-mer lookup within a list of references. Specifically, the core components of the index are (1) a minimal perfect hash function (MPHF), (2) a unitig sequence vector, (3) a unitig-to-reference table, and (4) a vector storing the position associated with each *k*-mer in the unitig sequence vector. The unitig sequence vector contains all the unitigs in the ccdBg. The Pufferfish index admits efficient exact search for *k*-mers, as well as longer matches that are unique in both the query string and colored compacted de Bruijn graph. These matches, called uni-MEM, were originally defined in deBGA^13^. A uni-MEM is a Maximal Exact Match (MEM) between the query sequence and a unitig. Using the combination of the MPHF and the position vector, a *k*-mer is mapped to a unitig in the unitig sequence vector. The *k*-mer is then extended to a uni-MEM via a linear scan of the query sequence and the unitig sequence vector. Each uni-MEM can appear in multiple different references, and since uni-MEMs must be completely contained within a unitig, it is possible for multiple uni-MEMs to be directly adjacent on both the query and some references where the unitig appears.

#### uni-MEM collection

The first step in read alignment is to collect exact matches shared between the query (single-end or paired-end reads) and the reference. In PuffAligner, this is accomplished by collecting the set of uni-MEMs that co-occur between the query and reference. PuffAligner starts processing the read from the left-end and looks up each *k*-mer that is encountered until a match to the index is found. Once a match is discovered, it is extended in both query and the reference until one of these termination conditions occur: (1) a mismatch is encountered, (2) the end of the query is reached, or (3) the end of the unitig is reached. This process results in a uni-MEM match shared between the query and reference. uni-MEMs where extension is terminated as a result of reaching the end of a unitig must later be examined and potentially “collpased” together to form MEMs with respect to the references on which they appear. If the uni-MEM extension is not terminated as a result of reaching the end of the query, then the position in the read is incremented by a small value and the same procedure is repeated for the next *k*-mer on the read. This process continues until either the uni-MEM extension terminates because the end of the query is reached, or because the last *k*-mer of the query is searched in the index. Here, we recall an important property of uni-MEM extension that is different from e.g. MEM extension or maximum mappable prefix (MMP) extension^5^. Due to the definition of the ccdBg, it is guaranteed that any *k*-mer appearing within a uni-MEM cannot appear in any other unintig in the ccdBg. Thus, extending *k*-mers to maximal uni-MEMs is, in some sense, safe with respect to greedy extension, as such extension will never cause missing a *k*-mer that would lead to another distinct uni-MEM shared between the query and reference. The concept of safe extension of kmer matches was introduced in^28^.

#### Filtering highly-repetitive uni-MEMs

In order to avoid expending computation on performing the subsequent steps on regions of reads mapping to highly-repeated regions of the reference, any uni-MEM that appears more than a user-defined number of times in the reference is discarded. In this manuscript, we use the threshold of 1000. This filter has a strong impact on the performance, since, even if one *k*-mer from the read maps to a highly-repetitive region of the reference, the following expensive steps of the alignment procedure should be performed for every mapping position of the uni-MEM to find the right alignment for the read, while the less repetitive uni-MEMs also map to the true origin of the read on the reference as well. The drawback of this filter is that for a very small fraction of the reads which are truly originating from a highly-repetitive region, all of the matched uni-MEMs will be filtered out and no hit remains for aligning the read. However, we find that in the case of aligning paired-end reads, usually one end of the read maps to a non-repetitive region, then, the alignment of the other end can be recovered using orphan recovery (explained in Section 4.4). Futheremore, we also provide a flag *–allowHighMultiMappers* that mitigates the effect of this filter for a slight tradeoff on the alignment performance.

#### uni-MEM compaction

For paired-end reads, PuffAligner aligns each end the read pairs individually. For each end, all the uni-MEMs are sorted on the basis of their positions on the reference. Consecutive uni-MEMs with no gap (both on the reference and the read) are merged into larger MEMs. The compactable uni-MEMs result from terminating the extension process due to reaching the end of a unitig. Such consecutive uni-MEMs can be safely compacted to form longer MEMs that will be used later in the MEM chaining algorithm. After the compaction of uni-MEMs, there is a list of MEMs which are shared sequences between the query and a set of reference positions, that are sorted based on the reference positions.

### 4.2 Finding promising MEM chains

As shown in figure 6, having all the MEMs (maximal exact matches) from a read to each target reference, the goal of this step is to find promising chains of MEMs that cover the most unique bases in the read in a concordant fashion and that can potentially lead to a high quality alignment.

**Figure 6:**
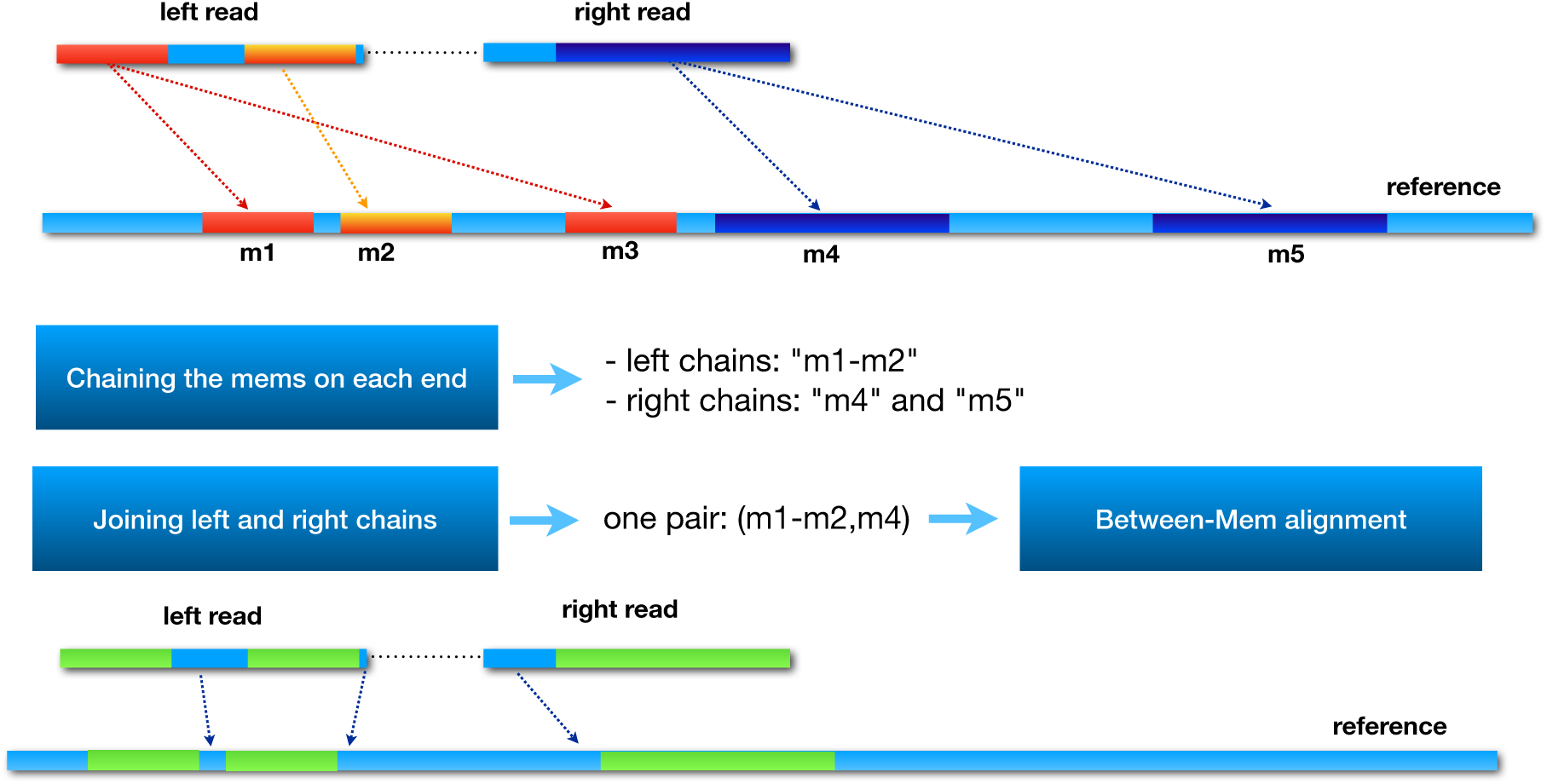
This figure shows the main steps of chaining and between-MEM alignment in the PuffAligner procedure via an example. In this example, m1, m2 and m3 are the projected MEMs from the left end of the read to the reference and m4 and m5 are the projected MEMs from the right end of the read. In the first step, the chaining algorithm chooses the best chain of MEMs that provide the highest coverage score for each end of the read, that is the m1-m2 chain for the left end and two single MEM chain for the right end. Then, the selected chains from each end are joined together to find the concordant pairs of chains, that is the (m1-m2, m4) pair for this read as m5 is too far from m1-m2. Then, the chain from each end will go through to the next step, between-MEM alignment. For the green areas (MEMs) no alignment is recalculated as they are exact matches. Only the un-matched blue parts of the chains (those nucleotides not occurring within a MEM) are aligned using a modified version of KSW2.

To accomplish this, we adopt the dynamic programming approach used in minimap2^11^ for finding co-linear chains of MEMs that are likely candidates to support high-scoring read alignments. As mentioned in minimap2, all the MEMs from a read *r* to the reference *t*, are sorted by the ending position of the MEMs on the reference. Then, this algorithm computes a score for each set of MEMs based on the number of unique covered bases in the read, the coverage score is also penalized by the length of the gaps, both in the read and reference sequence, between each consecutive pair of MEMs.

In PuffAligner, if the distance between two MEMs, *m*_1_ and *m*_2_, on the read and the reference is *d*_*r*_ and *d*_*t*_ respectively, these two MEMs should not be chained together if |*d*_*r*_ − *d*_*t*_| >*C*, where *C* is the maximum allowed gap. So, the penalization term, the *β* value in ^11^, in the coverage score computation is modified accordingly to prevent pairing of such MEMs.

Also, unlike what is done in minimap2^11^, rather than considering together the MEMs that are discovered on both ends of a paired-end read, we consider the chaining and chain filtering for each end of the read separately. This is done in order to make it easier to enforce the orientation consistency of the individual chains. Specifically, the chaining algorithm that is presented in^11^ introduces a transition in the recursion that can be used to switch between the MEMs that are part of one read and those that are part of the other. However, such switching makes it difficult to enforce the orientation consistency of the chains that are being built for each end of the read. One solution to this problem is to add another dimension to the dynamic programming table, encoding if one has already switched from the MEMs of one read end to the other, and the recurrence can be modified to allow only one switch from the one read end to the other, allowing enforcement of orientation consistency. However, we found that, in practice, simply chaining the read ends separately led to better performance.

Finally, we also adopt the heuristic proposed by minimap2^11^ when calculating the highest scoring chains. That is, when a MEM is added to the end of an existing chain, it is unlikely that a higher score for a chain containing this MEM will be obtained by adding it to a preceding chain. Thus, we consider only a small fixed number of rounds (by default 2) of preceding chains once we have found the first chain to which we can add the current MEM.

The chaining algorithm described above finds the best chains of MEMs shared between the read *r* and the reference *t* in orientation *o*. A chain is accepted if its score is greater than a configurable fraction, which we call the *consensusFraction*, times the maximum coverage score found for the read *r* to *any* reference. Throughout all the experiments in this manuscript the *consensusFraction* is set to 0.65. If a chain passes the consensus fraction threshold, we call it a *valid* chain. Additionally, rather than keeping all valid chains, we also filter highly-suboptimal chains with respect to the highest scoring chain *per-reference*. All valid chains shared between *r* and *t* are sorted by their scores, and chains having scores within 10% of the highest scoring chain for reference *t* are selected as potential mappings of the read *r* to the reference *t*. While these filters are essential for improving the throughput of the algorithm in finding the right alignment, they are carefully selected to have very little effect on the sensitivity of PuffAligner. For all the experiments in this manuscript, the same default settings of these parameters are used if not mentioned otherwise.

### 4.3 Computing base-to-base alignments between MEMs

After finding the high-scoring MEM chains for each reference sequence, a base-to-base alignment of the read to each of the candidate reference sequences is computed. Each selected chain implies a position on the reference sequence where the read might exhibit a high quality alignment. Thus, we can attempt to compute an optimal alignment of the read to the reference at this implied position, potentially allowing a small bit of padding on each side of the read. This approach utilizes the positional information provided by the MEM chains. However, the starting position of the alignments is not the only piece of information embedded in the chains. Rather each chain of MEMs consists of sub-sequences of the read (of size at least *k*, though often longer) which match exactly to the reference. While the optimal alignment of the read to the reference at the position being considered is not *guaranteed* to contain these exact matches as alignments of the corresponding substrings, this is almost always the case.

In PuffAligner, we aim to exploit the information from the long matches to accelerate the computation of the alignments. In fact, since only chains with relatively high coverage score are selected, a large portion of the read sequences are typically already matched to the positions in the reference with which they will be matched in the final optimal alignment. For instance, in fig. 6, for the final chains selected on the reference sequence, it is already known for the light blue, dark blue and green sub-sequences on the left end of the read precisely where they should align to the reference. Likewise this is the case for the yellow and purple sub-sequences on the right read. The unmapped regions of the reads are either bordered by the exact matches on both sides, or they occur at the either ends of the read sequence. PuffAligner skips aligning the whole read sequence by considering the exact matches of the MEMs to be part of the alignment solution. As a result, it is only required to compute the alignment of the small unmapped regions, which reduces the computation burden of the alignments.

When applying such an approach, two different types of alignment problems are introduced, which we call bounded sub-sequence alignment and ending sub-sequence alignment. For bounded sub-sequence alignment, we need to *globally* align some interval *i*_*r*_ of the read to an interval *i*_*t*_ of the reference. If *i*_*r*_ and *i*_*t*_ are of different lengths, the alignment solution will necessarily include insertions or deletions. If *i*_*r*_ and *i*_*t*_ are of the same length, then the optimal global alignment between them may or may not include indels. For each such bounded sub-sequence alignment, we determine the optimal alignment of *i*_*r*_ to *i*_*t*_ by computing a global pair-wise alignment between the intervals, and stitching the resulting alignment together with the exact matches that bound these regions.

Gaps at the beginning or the end of the read are symmetric cases, and so we describe, without loss of generality, the case where there is an unaligned interval of the read after the last MEM shared between the read and the reference. In this case, we need to solve the ending sub-sequence alignment problem. Here, the unaligned interval of the read consists of the substring spanning from the last nucleotide of the terminal MEM in the chain, up through the last nucleotide of the read. There is not a clearly-defined interval on the reference sequence. While the left end of the relevant reference interval is defined by the last reference nucleotide that is part of the bounding MEM, the right end of the reference interval should be determined by actually solving an extension or “end-free” alignment problem. We address this by performing extension alignment of the unaligned interval of the read to an interval of the reference that begins on the reference at the end of the terminal MEM, and extends for the length of the unaligned query interval plus the length of some problem-dependent buffer (which is determined by the maximum length difference between the read and reference intervals that would still admit an alignment within the acceptable score threshold).

An example of both of these cases is displayed in Figure 6. Specifically, an alignment of the read could be obtained by only solving two smaller alignment problems; one is the ending sub-sequence alignment of the unmapped region after the green MEM on the left read and the other is the bounded sub-sequence alignment of region on the right read bordered by the yellow and purple MEMs.

PuffAligner uses KSW2^11,46^ for computing the alignments of the gaps between the MEMs and for aligning the ending sequences. KSW2 exposes a number of alignment modes such as global and extension alignments. For aligning the bounded regions, KSW2 alignment in the global mode is performed, and for the gaps at the beginning or end of reads, PuffAligner uses the extension mode to find the best possible alignment of that region. PuffAligner, by default, uses a match score of 2 and mismatch penalty of 4. For indels, PuffAligner uses an affine gap scoring schema with gap open penalty of 5 and gap extension penalty of 3. In PuffAligner, after computing the alignment score for each read, only the alignments with a score higher than *τ* times the maximum possible score for the read are reported. The value of *τ* is controlled by the option *–minScoreFraction*, which is set to 0.65 by default.

#### 4.3.1 Enhancing alignment computation

By only aligning the read’s sub-sequences that are not included in the MEMs, the size of alignment problems being solved in PuffAligner are often much shorter than the length of the read. However, to further speed up alignment, we also incorporate a number of other techniques to improve the performance of the alignment calculation. We describe the most important of these below:

- **Skipping alignment calculation by recognizing perfect chains and alignment caching:** It is possible to avoid the alignment computation completely in a considerable number of cases. In fact, as has been explained in previous work^28^, the alignment calculation step can be completely skipped if the set of exact matches for each chain covers the whole read. PuffAligner skips alignment for cases where the coverage score of chains of MEMs is the length of the read, and assigns a total matched CIGAR string for that alignment. Alignment computation of a read might be also skipped if the same alignment problem has been already detected and computed for this read. For example, in the case of RNA seq data, reads often map to the same exons on different transcripts. In such cases, each alignment solution for a read is stored in a cache (a hash table) so that if the same alignment problem is detected, the solution can be directly retrieved from the cache, and no further computation is required (see supplementary table S1).
- **Early stopping of the alignment computation when a valid score cannot be achieved:** While care is taken to produce only high-scoring chains between the read and reference, it is nonetheless the case that the majority of the chains do not lead to an alignment of acceptable quality. Since the minimum acceptable alignment score is immediately known based on *τ* and the length of the read, the base-to-base alignment calculation can be terminated at any point where it becomes imposible for the minimum required alignment score to be obtained. This approach can be applied both during the KSW2 alignment calculation, and also after the alignment calculation of each gap is completed. During this procedure, for each base at position *i*, starting from position 1 on the read of length *n*, if the best alignment score *p* up to the *i*-th position is *s*_*i*_, we can calculate the maximum possible alignment score, *s*_*max*_, that might be achieved starting at this location given the current alignment score by:

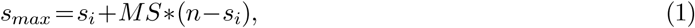

where *MS* is the score assigned to each match. If *s*_*max*_ is smaller than minimum required score for accepting the alignment, the alignment calculation can be immediately terminated, since it is already known that this anchor is not going to yield a valid alignment for this read.
- **Full-sensitivity banded alignment:** KSW2 is able to perform banded alignment to make alignment calculation more efficient. In this mode, the dynamic programming matrix for the alignment problem is only filled out along the sub-diagonals out to a certain distance *d* away from the main diagonal. If one is guaranteed that any valid alignment must have fewer than *d* insertions or deletions, then the alignment must not exit these bands of the dynamic programming matrix. Note that alignments with >*d* indels can be represented within these bands as insertions and deletions move in opposite anti-diagonal directions, but it is certainly the case that no alignment with ≤*d* indels can exit these bands. By calculating the maximum number of gaps (insertions or deletions) allowed in each sub-alignment probem, in a way that we are certain that any alignment having greater than this number of gaps must drop below the acceptable threshold, we utilize the banded alignment in KSW2 within each sub-alignment problem without losing any sensitivity with respect to non-banded alignment.

### 4.4 Joining mappings for read ends and orphan recovery

Finally, once alignments have been computed for the individual ends of a read, they must be paired together to produce valid alignments for the entire fragment. At this point in the process, on each reference sequence, there are a number of locations where the left end of each read or the right end of each read, or both, are mapped to the reference. For the purpose of determining which mappings will be reported as a valid pair, the mappings are joined together only if they occur on opposite strands of the reference, and if they are within a maximum allowed fragment length. There are two different types of paired-end alignments that can be reported by PuffAligner; concordant and discordant. If PuffAligner is disallowed from reporting discordant alignments, then the mapping orientation of the left and right end should agree with the library preparation protocols of the reads. PuffAligner first tries to find concordant mapping pairs on a reference sequence, and if no concordant mapping is discovered and the tool is being run in a mode where discordant mappings are allowed, then PuffAligner reports pairs that map discordantly. Here, discordant pairs may be pairs that do not, for example, obey the requirement of originating from opposite strands. While this is not expected to happen frequently, it may occur if there has been an inversion in the sequenced genome with respect to the reference.

#### Orphan recovery

If there is no valid paired-end alignment for a fragment (either concordant or discordant, if the latter is allowed), then PuffAligner will attempt to perform orphan recovery. The term “orphan” refers to one end of paired-end read that is confidently aligned to some genomic position, but for which the other read end is not aligned nearby (and paired). To perform orphan recovery, PuffAligner examines the reference sequence downstream of the mapped read (or upstream if the mapped read is aligned to the reverse complement strand) and directly performs dynamic programming to look for a valid mapping of the unmapped read end. For this purpose, we use the “fitting” alignment functionality of edlib^47^ to perform a simple Levenshtein distance based alignment that will subsequently be re-scored by KSW2. Finally, if, after attempting orphan recovery, there is still no valid paired-end mapping for the fragment, then orphan alignments are reported by PuffAligner (unless the “`--noOrphans`” flag is passed).

## Competing interests

RP is a co-founder of Ocean Genomics Inc.

## Acknowledgements

The authors would like to thank Laraib Iqbal Malik for her help and suggestions for improving the manuscript.

## Funding

This work has been funded by R01 HG009937, NSF CCF-1750472, and NSF CNS-1763680 to R.P. The funders had no role in study design, data collection and analysis, decision to publish, or preparation of the manuscript.

## Supplementary Material for

**Table S1:** The percentage of aligner engine calls skipped in the alignment calculation pipeline.

| sample | Cache Hits | Perfect Chains | None Alignable | Total Skipped |
| --- | --- | --- | --- | --- |
| DNA-seq experimental | 52.89% | 19.01% | 0.71% | 72.67% |
| RNA-seq simulated | 28.69% | 50.80% | 0.97% | 80.46% |
| Metagenomic simulated | 61.10% | 31.33% | 0.00 % | 92.43% |

**Figure S1:**
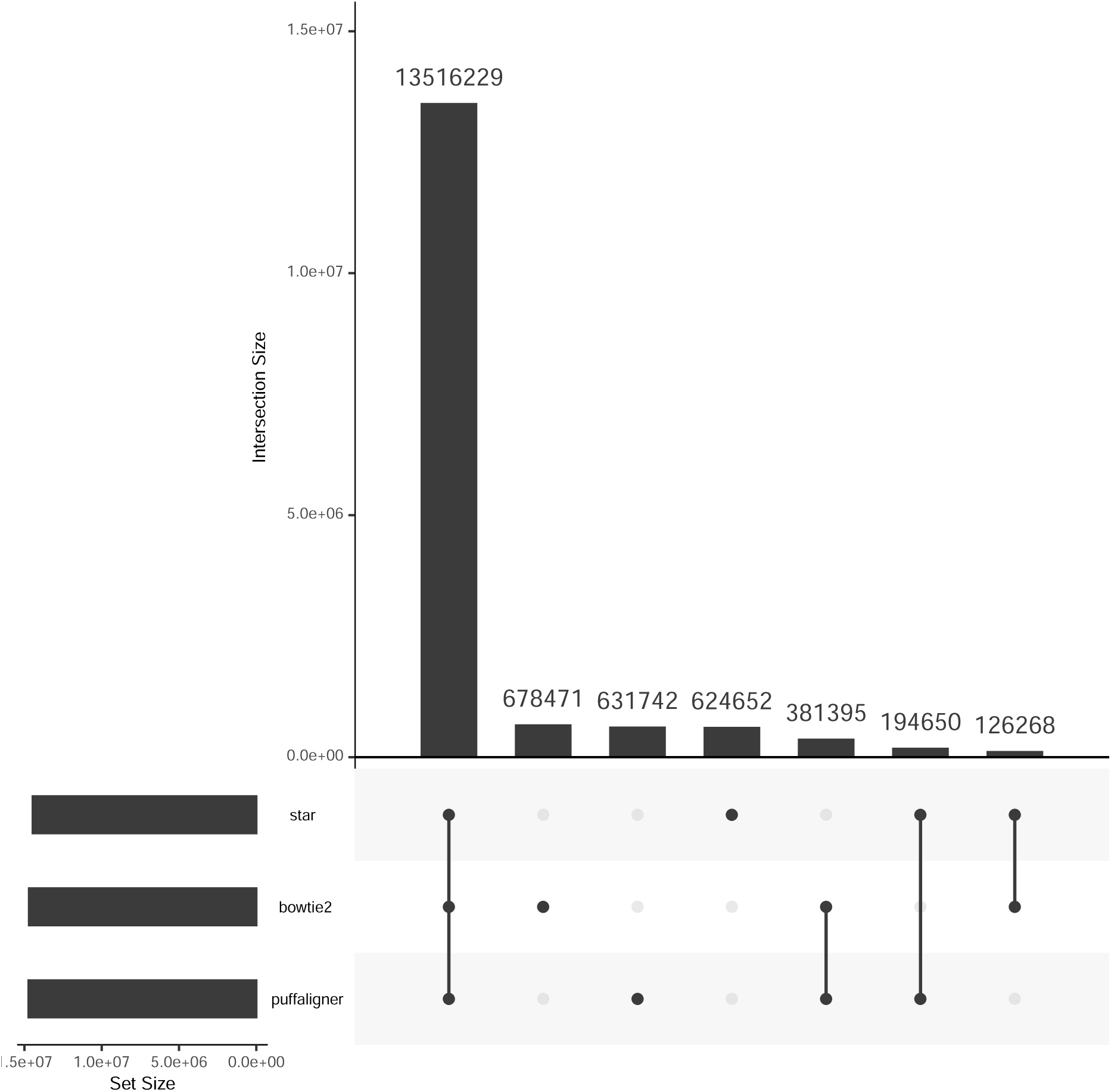
Upset plot showing the agreement of the alignments found by different tools based on the location of the mappings

**Table S2:**
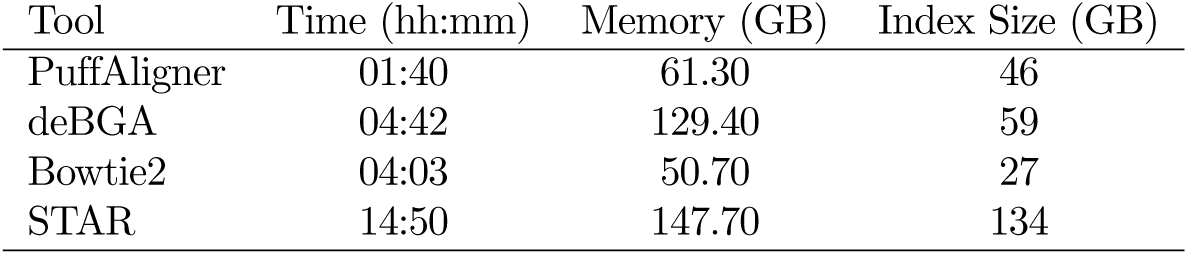
The construction benchmark and final index size for each of the tools over 4000 selected bacterial genomes.

| Tool | Time (hh:mm) | Memory (GB) | Index Size (GB) |
| --- | --- | --- | --- |
| PuffAligner | 01:40 | 61.30 | 46 |
| deBGA | 04:42 | 129.40 | 59 |
| Bowtie2 | 04:03 | 50.70 | 27 |
| STAR | 14:50 | 147.70 | 134 |

**Table S3:** Basic information for samples selected for simulating mock bulk metagenomic samples.

| Accession | Project description | # of reads | # of reads aligned to 4k selected reference | # of simulated reads | # of references of origin for the simulated reads |
| --- | --- | --- | --- | --- | --- |
| SRR10948222 | Finding sub-biocrust soil microbial communities in Mojave Desert, California, United States | 27,296,270 | 200k | 5,550,650 | 98 |
| SRR11283975 | The impact of different acidification degrees on the bacterial community in the Jiaodong Peninsula, China | 35.5k | 8,333 | 1,012,176 | 92 |
| SRR11496426 | The composition, genetics characteristics and structure of the microbial communities of the oil site of Uzon Caldera | 42.3k | 30,203 | 1,029,382 | 179 |

The formula for calculating the metrics used for evaluating the abundance estimation results in the manuscript are as follows. The metrics are Mean Absolute Relative Difference (MARD), Mean Absolute Error (MAE), and Mean Squared Log Error (MSLE).

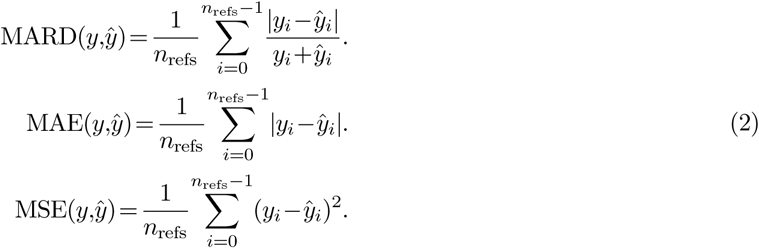

**Table S4:** Alignment Distribution for two samples of 500,000 simulated reads from reference sequences NC_004718.3 (known as SARS Coronavirus) and NC_014470.1 (Bat Coronavirus assembled in 2008) respectively.

| Sample's Origin | Alignment Mode | Tool | NC_045512.2 (Covid19) | NC_004718.3 (SARS Coronavirus) | NC_014470.1 (Bat Coronavirus) | Other Assembled References |
| --- | --- | --- | --- | --- | --- | --- |
| SARS | Primary | PuffAligner | 0 | 500,000 | 0 | 0 |
|  |  | Bowtie2 | 24 | 499,975 | 0 | 0 |
|  |  | STAR | 0 | 499,998 | 0 | 0 |
|  |  | deBGA | 0 | 499,995 | 0 | 5 |
|  | Up to 20 | PuffAligner | 116 | 500,000 | 486 | 0 |
|  |  | Bowtie2 | 21,546 | 499,999 | 7,205 | 0 |
|  |  | STAR | 0 | 499,998 | 0 | 0 |
|  |  | deBGA | 0 | 499,995 | 0 | 5 |
|  | Up to 200 | PuffAligner | 116 | 500,000 | 486 | 0 |
|  |  | Bowtie2 | 21,546 | 499,999 | 7,205 | 0 |
|  |  | STAR | 0 | 499,998 | 0 | 0 |
|  |  | deBGA | 0 | 499,995 | 0 | 5 |
|  | Best strata | PuffAligner | 0 | 500,000 | 0 | 0 |
|  |  | STAR | 0 | 499,998 | 0 | 0 |
| Bat2008 | Primary | PuffAligner | 0 | 0 | 500,000 | 0 |
|  |  | Bowtie2 | 0 | 0 | 499,999 | 0 |
|  |  | STAR | 0 | 0 | 499,999 | 0 |
|  |  | deBGA | 0 | 0 | 499,991 | 9 |
|  | Up to 20 | PuffAligner | 32 | 494 | 500,000 | 0 |
|  |  | Bowtie2 | 2,343 | 7,127 | 499,999 | 0 |
|  |  | STAR | 0 | 0 | 499,999 | 0 |
|  |  | deBGA | 0 | 0 | 499,991 | 0 |
|  | Up to 200 | PuffAligner | 32 | 494 | 500,000 | 0 |
|  |  | Bowtie2 | 2,343 | 7,127 | 499,999 | 0 |
|  |  | STAR | 0 | 0 | 499,999 | 0 |
|  |  | deBGA | 0 | 0 | 499,991 | 9 |
|  | Best strata | PuffAligner | 0 | 0 | 500,000 | 0 |
|  |  | STAR | 0 | 0 | 499,999 | 0 |

**Table S5:** Alignment accuracy of the tools over different accuracy metrics for the mock sample simulated from real samples with accession IDs SRR11283975 and SRR11496426.

| Accession ID | Alignment Mode | Tool | Spearman | MARD | MAE | MSLE |
| --- | --- | --- | --- | --- | --- | --- |
| SRR11283975<br>Jiandong Peninsula | Primary | PuffAligner | 0.71 | 0.024 | 0.43 | 0.04 |
|  |  | Bowtie2 | 0.615 | 0.04 | 0.64 | 0.07 |
|  |  | STAR | 0.727 | 0.02 | 0.406 | 0.04 |
|  |  | deBGA | 0.274 | 0.521 | 106.78 | 3.79 |
|  | Up to 20 | PuffAligner | 0.942 | 0.003 | 0.07 | 0.00 |
|  |  | Bowtie2 | 0.909 | 0.005 | 0.05 | 0.00 |
|  |  | STAR | 0.946 | 0.003 | 0.087 | 0.00 |
|  |  | deBGA | 0.277 | 0.489 | 101.39 | 3.37 |
|  | Up to 200 | PuffAligner | 0.979 | 0.001 | 0.07 | 0 |
|  |  | Bowtie2 | 0.97 | 0.002 | 0.04 | 0.00 |
|  |  | STAR | 0.951 | 0.003 | 0.086 | 0.00 |
|  |  | deBGA | 0.278 | 0.483 | 100.96 | 3.29 |
|  | Best strata | PuffAligner | 0.979 | 0.001 | 0.063 | 0 |
|  |  | STAR | 0.951 | 0.003 | 0.086 | 0.00 |
| SRR11496426<br>Uzon Caldera | Primary | PuffAligner | 0.568 | 0.112 | 32.55 | 0.95 |
|  |  | Bowtie2 | 0.53 | 0.14 | 38.06 | 1.10 |
|  |  | STAR | 0.559 | 0.118 | 31.823 | 0.83 |
|  |  | deBGA | 0.367 | 0.566 | 115.88 | 3.57 |
|  | Up to 20 | PuffAligner | 0.789 | 0.03 | 7.43 | 0.24 |
|  |  | Bowtie2 | 0.74 | 0.042 | 10.83 | 0.30 |
|  |  | STAR | 0.713 | 0.049 | 6.939 | 0.17 |
|  |  | deBGA | 0.368 | 0.554 | 109.29 | 3.32 |
|  | Up to 200 | PuffAligner | 0.865 | 0.017 | 5.64 | 0.11 |
|  |  | Bowtie2 | 0.879 | 0.015 | 7.21 | 0.13 |
|  |  | STAR | 0.724 | 0.045 | 6.496 | 0.13 |
|  |  | deBGA | 0.369 | 0.549 | 108.99 | 3.27 |
|  | Best strata | PuffAligner | 0.85 | 0.019 | 5.571 | 0.09 |
|  |  | STAR | 0.723 | 0.046 | 6.544 | 0.13 |

**Figure S2:**
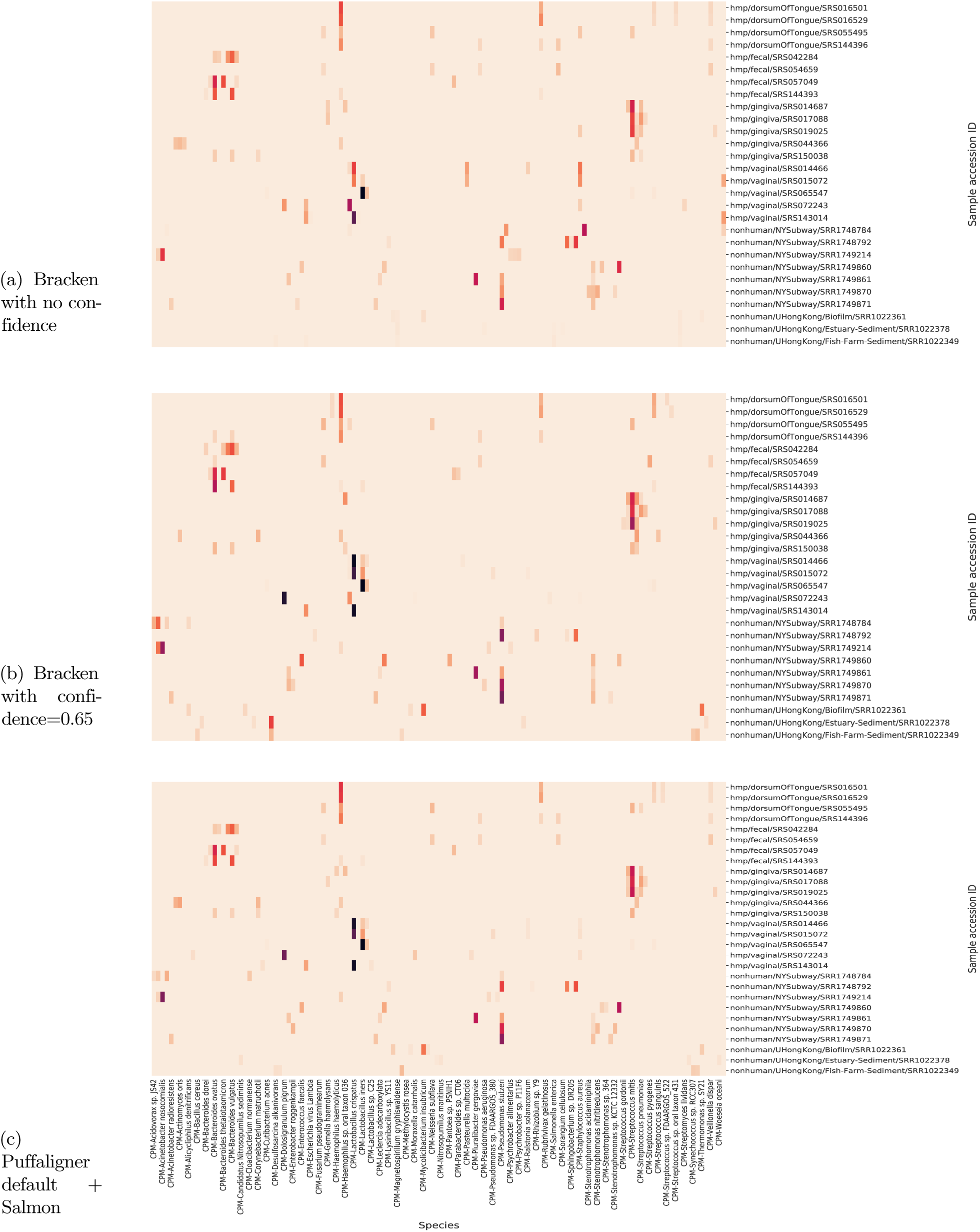
Heatmap showing 5 most popular species over 28 samples through three pipelines of Kraken2(no confidence)+Bracken, Kraken2(confidence=0.65)+Bracken and default Puffaligner+Salmon. Overall, we observe more similarity between Puffaligner and Bracken with confidence of 0.65. However, there are cases where applying the confidence filter to Kraken make the results diverge from Puffaligner pipeline for example “Screptococcus gorbanii” considered abundant in a subway sample in the first and third heatmap, whereas Kraken2(confidence=0.65) does not detect the microorganism as abundant.

**Figure S3:**
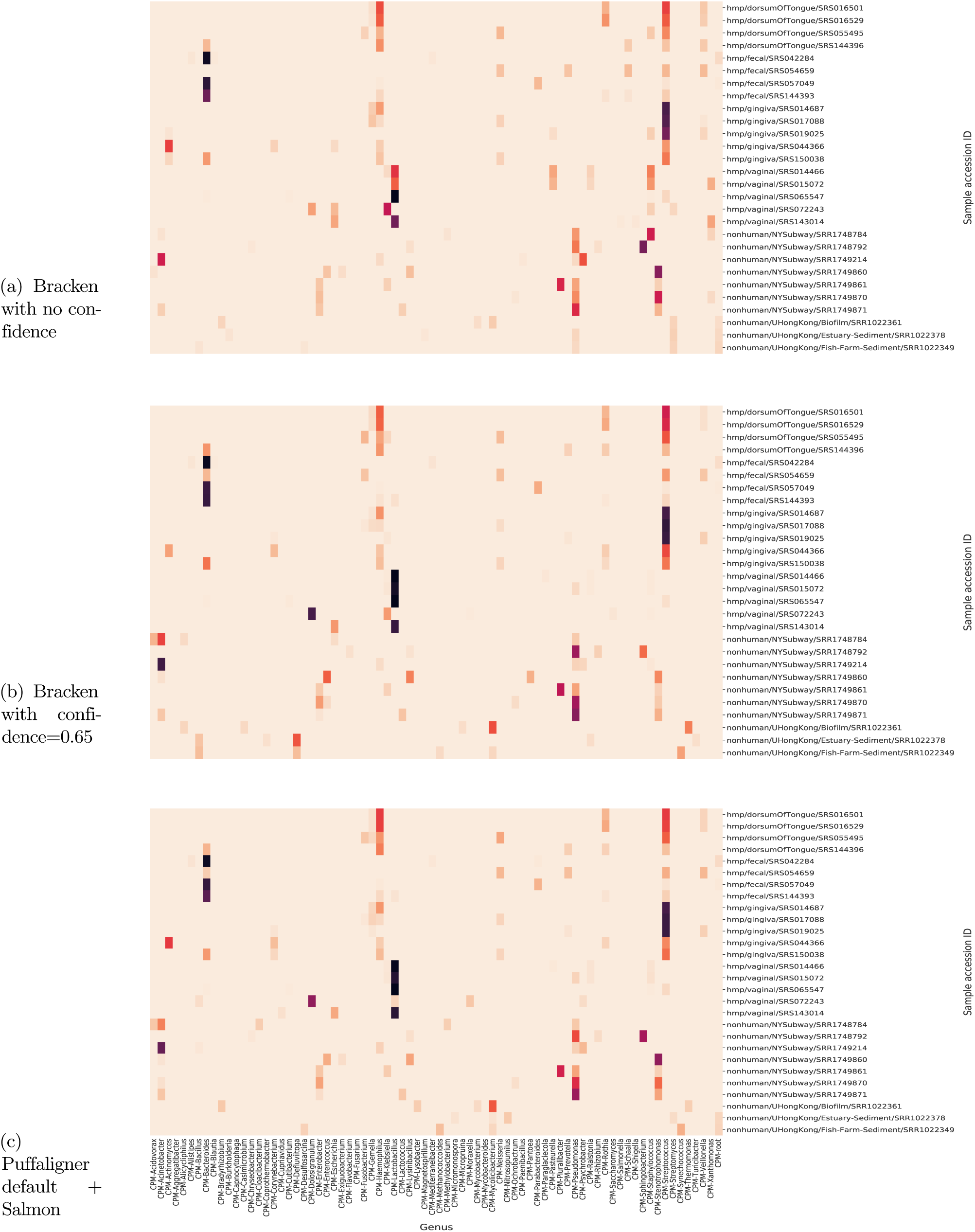
Heatmap showing 5 most popular genera over 28 samples through three pipelines of Kraken2(no confidence)+Bracken, Kraken2(confidence=0.65)+Bracken and default Puffaligner+Salmon.

## Footnotes

* https://www.internationalgenome.org/data-portal/sample/HG00190

† https://www.gencodegenes.org/human/release_33.html

‡ ftp://ftp.ensembl.org/pub/release-98/fasta/homo_sapiens/dna/

§ https://github.com/DerrickWood/kraken2/wiki/Manual

¶ https://github.com/DerrickWood/kraken2/issues/167

## Notes

### Competing Interest Statement

The authors have declared no competing interest.

## Bibliography

[1] Heng Li and Richard Durbin. Fast and accurate short read alignment with Burrows–Wheeler transform. Bioinformatics, 25(14):1754–1760, 2009.

[2] Ben Langmead and Steven L Salzberg. Fast gapped-read alignment with Bowtie 2. Nature Methods, 9(4): 357, 2012.

[3] Daehwan Kim, Ben Langmead, and Steven L Salzberg. Hisat: a fast spliced aligner with low memory requirements. Nature methods, 12(4):357–360, 2015.

[4] Daehwan Kim, Joseph M Paggi, Chanhee Park, Christopher Bennett, and Steven L Salzberg. Graph-based genome alignment and genotyping with hisat2 and hisat-genotype. Nature biotechnology, 37(8):907–915, 2019.

[5] Alexander Dobin, Carrie A Davis, Felix Schlesinger, Jorg Drenkow, Chris Zaleski, Sonali Jha, Philippe Batut, Mark Chaisson, and Thomas R Gingeras. STAR: ultrafast universal RNA-seq aligner. Bioinformatics, 29 (1):15–21, 2013.

[6] Yang Liao, Gordon K Smyth, and Wei Shi. The subread aligner: fast, accurate and scalable read mapping by seed-and-vote. Nucleic acids research, 41(10):e108–e108, 2013.

[7] Matei David, Misko Dzamba, Dan Lister, Lucian Ilie, and Michael Brudno. Shrimp2: sensitive yet practical short read mapping. Bioinformatics, 27(7):1011–1012, 2011.

[8] Can Alkan, Jeffrey M Kidd, Tomas Marques-Bonet, Gozde Aksay, Francesca Antonacci, Fereydoun Hormozdiari, Jacob O Kitzman, Carl Baker, Maika Malig, Onur Mutlu, et al. Personalized copy number and segmental duplication maps using next-generation sequencing. Nature genetics, 41(10):1061, 2009.

[9] Faraz Hach, Fereydoun Hormozdiari, Can Alkan, Farhad Hormozdiari, Inanc Birol, Evan E Eichler, and S Cenk Sahinalp. mrsfast: a cache-oblivious algorithm for short-read mapping. Nature methods, 7(8):576, 2010.

[10] Chirag Jain, Alexander Dilthey, Sergey Koren, Srinivas Aluru, and Adam M. Phillippy. A fast approximate algorithm for mapping long reads to large reference databases. Journal of Computational Biology, 25(7): 766–779, July 2018. doi:10.1089/cmb.2018.0036. URL https://doi.org/10.1089/cmb.2018.0036.

[11] Heng Li. Minimap2: pairwise alignment for nucleotide sequences. Bioinformatics, 34(18):3094–3100, 2018.

[12] Nicolas L Bray, Harold Pimentel, Páll Melsted, and Lior Pachter. Near-optimal probabilistic RNA-seq quantification. Nature Biotechnology, 34(5):525, 2016.

[13] Bo Liu, Hongzhe Guo, Michael Brudno, and Yadong Wang. debga: read alignment with de bruijn graph-based seed and extension. Bioinformatics, 32(21):3224–3232, 2016.

[14] Antoine Limasset, Bastien Cazaux, Eric Rivals, and Pierre Peterlongo. Read mapping on de bruijn graphs. BMC bioinformatics, 17(1):237, 2016.

[15] Mahdi Heydari, Giles Miclotte, Yves Van de Peer, and Jan Fostier. Browniealigner: accurate alignment of illumina sequencing data to de bruijn graphs. BMC bioinformatics, 19(1):311, 2018.

[16] Fatemeh Almodaresi, Hirak Sarkar, Avi Srivastava, and Rob Patro. A space and time-efficient index for the compacted colored de bruijn graph. Bioinformatics, 34(13):i169–i177, 2018.

[17] Zamin Iqbal, Mario Caccamo, Isaac Turner, Paul Flicek, and Gil McVean. De novo assembly and genotyping of variants using colored de bruijn graphs. Nature genetics, 44(2):226, 2012.

[18] Martin D Muggli, Alexander Bowe, Noelle R Noyes, Paul S Morley, Keith E Belk, Robert Raymond, Travis Gagie, Simon J Puglisi, and Christina Boucher. Succinct colored de bruijn graphs. Bioinformatics, 33(20): 3181–3187, 2017.

[19] Fatemeh Almodaresi, Prashant Pandey, and Rob Patro. Rainbowfish: A succinct colored de bruijn graph representation. In 17th International Workshop on Algorithms in Bioinformatics (WABI 2017). Schloss Dagstuhl-Leibniz-Zentrum fuer Informatik, 2017.

[20] Prashant Pandey, Fatemeh Almodaresi, Michael A Bender, Michael Ferdman, Rob Johnson, and Rob Patro. Mantis: A fast, small, and exact large-scale sequence-search index. Cell systems, 7(2):201–207, 2018.

[21] 1000 Genomes Project Consortium et al. A global reference for human genetic variation. Nature, 526(7571): 68–74, 2015.

[22] Shifu Chen, Yanqing Zhou, Yaru Chen, and Jia Gu. fastp: an ultra-fast all-in-one fastq preprocessor. Bioinformatics, 34(17):i884–i890, 2018.

[23] Adam Frankish, Mark Diekhans, Anne-Maud Ferreira, Rory Johnson, Irwin Jungreis, Jane Loveland, Jonathan M Mudge, Cristina Sisu, James Wright, Joel Armstrong, et al. Gencode reference annotation for the human and mouse genomes. Nucleic acids research, 47(D1):D766–D773, 2019.

[24] Jake R Conway, Alexander Lex, and Nils Gehlenborg. Upsetr: an r package for the visualization of intersecting sets and their properties. Bioinformatics, 33(18):2938–2940, 2017.

[25] Avi Srivastava, Laraib Malik, Hirak Sarkar, Mohsen Zakeri, Fatemeh Almodaresi, Charlotte Soneson, Michael I Love, Carl Kingsford, and Rob Patro. Alignment and mapping methodology influence transcript abundance estimation. BioRxiv, page 657874, 2019.

[26] Manuel Holtgrewe. Mason: a read simulator for second generation sequencing data. 2010.

[27] Rob Patro, Geet Duggal, Michael I Love, Rafael A Irizarry, and Carl Kingsford. Salmon provides fast and bias-aware quantification of transcript expression. Nature Methods, 14(4):417, 2017.

[28] Hirak Sarkar, Mohsen Zakeri, Laraib Malik, and Rob Patro. Towards selective-alignment: Bridging the accuracy gap between alignment-based and alignment-free transcript quantification. In Proceedings of the 2018 ACM International Conference on Bioinformatics, Computational Biology, and Health Informatics, pages 27–36, Washington DC, USA, 2018. ACM. URL http://doi.acm.org/10.1145/3233547.3233589.

[29] Hy Vuong, Thao Truong, Thang Tran, and Son Pham. A revisit of rsem generative model and its em algorithm for quantifying transcript abundances. bioRxiv, page 503672, 2018.

[30] Alyssa C Frazee, Andrew E Jaffe, Ben Langmead, and Jeffrey T Leek. Polyester: simulating rna-seq datasets with differential transcript expression. Bioinformatics, 31(17):2778–2784, 2015.

[31] John Lonsdale, Jeffrey Thomas, Mike Salvatore, Rebecca Phillips, Edmund Lo, Saboor Shad, Richard Hasz, Gary Walters, Fernando Garcia, Nancy Young, et al. The genotype-tissue expression (gtex) project. Nature genetics, 45(6):580, 2013.

[32] Bo Li and Colin N Dewey. RSEM: accurate transcript quantification from RNA-Seq data with or without a reference genome. BMC Bioinformatics, 12(1):323, 2011.

[33] Ben Langmead. Aligning short sequencing reads with bowtie. Current protocols in bioinformatics, 32(1): 11–7, 2010.

[34] Fan Wu, Su Zhao, Bin Yu, Yan-Mei Chen, Wen Wang, Zhi-Gang Song, Yi Hu, Zhao-Wu Tao, Jun-Hua Tian, Yuan-Yuan Pei, et al. A new coronavirus associated with human respiratory disease in china. Nature, 579 (7798):265–269, 2020.

[35] Pavel V Baranov, Clark M Henderson, Christine B Anderson, Raymond F Gesteland, John F Atkins, and Michael T Howard. Programmed ribosomal frameshifting in decoding the sars-cov genome. Virology, 332 (2):498–510, 2005.

[36] Yixuan Wang, Yuyi Wang, Yan Chen, and Qingsong Qin. Unique epidemiological and clinical features of the emerging 2019 novel coronavirus pneumonia (covid-19) implicate special control measures. Journal of medical virology, 92(6):568–576, 2020.

[37] Tao Zhang, Qunfu Wu, and Zhigang Zhang. Probable pangolin origin of sars-cov-2 associated with the covid-19 outbreak. Current Biology, 2020.

[38] Xiaolu Tang, Changcheng Wu, Xiang Li, Yuhe Song, Xinmin Yao, Xinkai Wu, Yuange Duan, Hong Zhang, Yirong Wang, Zhaohui Qian, et al. On the origin and continuing evolution of sars-cov-2. National Science Review, 2020.

[39] PI: Kirsten Fisher. Sub-biocrust soil microbial communities from mojave desert, california, united states - 8hms. Sequence Read Archive (SRA) [Internet]. Bethesda (MD): National Library of Medicine (US), National Center for Biotechnology Information; 2009, 1 2020. submitted to JGI at 2019-09-20; Available from: https://www.ncbi.nlm.nih.gov/sra/.

[40] Jennifer Lu, Florian P Breitwieser, Peter Thielen, and Steven L Salzberg. Bracken: estimating species abundance in metagenomics data. PeerJ Computer Science, 3:e104, 2017.

[41] Mark Reppell and John Novembre. Using pseudoalignment and base quality to accurately quantify microbial community composition. PLoS computational biology, 14(4):e1006096, 2018.

[42] Scott Federhen. The ncbi taxonomy database. Nucleic acids research, 40(D1):D136–D143, 2012.

[43] MetaSUB International Consortium et al. The metagenomics and metadesign of the subways and urban biomes (metasub) international consortium inaugural meeting report, 2016.

[44] Liping Ma, Bing Li, and Tong Zhang. Abundant rifampin resistance genes and significant correlations of antibiotic resistance genes and plasmids in various environments revealed by metagenomic analysis. Applied microbiology and biotechnology, 98(11):5195–5204, 2014.

[45] Dirk Gevers, Rob Knight, Joseph F Petrosino, Katherine Huang, Amy L McGuire, Bruce W Birren, Karen E Nelson, Owen White, Barbara A Methé, and Curtis Huttenhower. The human microbiome project: a community resource for the healthy human microbiome. PLoS Biol, 10(8):e1001377, 2012.

[46] Hajime Suzuki and Masahiro Kasahara. Introducing difference recurrence relations for faster semi-global alignment of long sequences. BMC Bioinformatics, 19(1):45, 2018.

[47] Martin Šošić and Mile Šikić. Edlib: a C/C++ library for fast, exact sequence alignment using edit distance. Bioinformatics, 33(9):1394–1395, 2017.

